# How do brassinosteroids activate their receptors?

**DOI:** 10.1101/630640

**Authors:** Alexander S. Moffett, Diwakar Shukla

**Affiliations:** Center for Biophysics and Quantitative Biology, University of Illinois at Urbana-Champaign; Department of Chemical and Biomolecular Engineering, University of Illinois at Urbana-Champaign; Department of Plant Biology, University of Illinois at Urbana-Champaign; Beckman Institute for Advanced Science and Technology, University of Illinois at Urbana-Champaign

**Keywords:** molecular dynamics, plant hormone, computational biology, signal transduction, protein structure

## Abstract

Brassinosteroids (BRs) are an important class of plant growth hormones which signal through BRI1 and BAK1, leucine-rich repeat receptor-like kinases (LRR-RLKs). When bound to the BRI1 island domain, BRs act as a “molecular glue”, mediating interactions between BRI1 and BAK1 extracellular domains. However, it is unclear how much other factors contribute to BR-induced BRI1-BAK1 association, including stabilization of the BRI1 island domain and large conformational changes in BRI1. We use several molecular dynamics simulation-based methods to explore the contributions of each mechanism to BR-dependent *Arabidopsis thaliana* BRI1-BAK1 association. We find that specific BR interactions make major contributions to BRI1-BAK1 association free energy. BR binding stabilizes the BRI1 island domain, while BRI1 undergoes a large conformational change to form a secondary interface with BAK1. These results suggest that each mechanism plays a part in BR signal transduction while raising questions about the functional role of conformational dynamics in other LRR-RLKs.

## 1 Introduction

Brassinosteroids (BRs) are a class of plant hormones involved in a number of growth and developmental processes [1]. One such BR, brassinolide (BL), binds to the ectodomain of the leucine-rich repeat receptor-like kinase (LRR-RLK) BRASSINOSTEROID-INSENSITIVE 1 (BRI1), stabilizing BRI1 interaction with its coreceptor BRI1-ASSOCIATED KINASE 1 (BAK1, also called SERK3)[2, 3, 4, 5, 6, 7], a versatile LRR-RLK playing a role in numerous signaling pathways [8, 9, 10]. After BL binds to BRI1, the intracellular kinase domains of BRI1 and BAK1 reciprocally phosphorylate one another, activating downstream signaling pathways and ultimately affecting gene expression [11].

An atomistic picture of brassinosteroid signaling initiation has begun to emerge from a number of structural and computational studies [2, 3, 5, 6, 7, 12, 13, 14, 15, 16, 17, 18, 19]. The predominant hypothesis concerning the mechanism of BL-induced BR1-BAK1 association is that BL acts as a “molecular glue”, mediating interactions between the BRI1 and BAK1 extracellular domains (ECDs), as observed in crystal structures[5, 6]. Several researchers have suggested that BL binding stabilizes the BRI1 island domain, which could reduce the entropic cost of BRI1-BAK1 association [2, 3, 20]. However, the exact molecular events that unfold upon binding of BL to BRI1 remain unclear, especially in light of evidence that BRI1 and BAK1 preform dimers *in vivo* in the absence of BL [21, 22]. While recent work has provided quantitative thermodynamic data concerning BL binding to BRI1 and BRI1-BAK1 association [19], the nanoscale details of how hormone binding causes the BRI1-BAK1 complex to assemble and activate remain largely unknown.

In addition to the molecular glue and BRI1 island domain stabilization hypotheses, the possibility of BL-induced shifts in the BRI1 conformational equilibrium, stabilizing the BRI1-BAK1 complex, has not been conclusively ruled out. In fact, it remains possible that all three proposed mechanisms play an important role in BRI1-BAK1 activation. The BRI1 island domain B-factors are lower in holo BRI1 crystal structures than in apo structures[2, 3], which suggests that BL binding to BRI1 reduces fluctuations in the BRI1 island domain, providing a stable platform for BAK1 interaction. In a BRI1-BAK1-BL crystal structure [6], the two BL hydroxyl groups of carbons 2 and 3 on the A steroid ring interact with the BAK1 backbone and H61 side chain, while the phenyl group of BAK1 F60 lies in a plane parallel to the BL steroid rings. *In vitro* experiments on the BRI1 and BAK1 ECD domains found that their BL-dependent association was also dependent on pH [6], suggesting that the protonation state of H61 may play an important role in controlling BRI1-BAK1 association. Although no previous evidence of large BRI1 conformational changes exists, the structure and dynamics of the BRI1 ECD has only been investigated in the context of low temperature crystal packing, and it remains unclear whether global conformational changes in the BRI1 ECD occur or if BL binding influences BRI1 conformational dynamics.

In this study, we used molecular dynamics (MD) simulations to examine at the atomic scale several proposed mechanisms as to how BL binding causes the association of BRI1 and BAK1. Starting from a crystal structure of the BRI1-BAK1-BL complex[6], we calculated the absolute apo and holo BRI1-BAK1 ECD association free energies for a truncated BRI1 ECD (tBRI1) containing LRRs 13-25 and the island domain. We found that with protonation states consistent with a pH 7 solution, BL binding only moderately stabilizes the tBRI1-BAK1 complex. By removing the N-terminal region of the BRI1 ECD, we quantified the impact of BL-mediated interactions on BRI1-BAK1 complex stability under the assumption that BRI1 LRRs 1-12 are static with respect to the rest of the BRI1 ECD and do not play any role in BRI1-BAK1 association. We used alchemical free energy calculations to explore the impact of pH on BL-induced tBRI1-BAK1 association, finding that double protonation of BAK1 H61 plays a central role in stabilizing the holo complex over the apo complex. We performed Gaussian accelerated molecular dynamics (GAMD) simulations of apo and holo tBRI1 in order to investigate the effect of BL binding on island domain dynamics. The tBRI1 island domain was stabilized by the presence of BL, while the apo island domain exhibited a greater degree of flexibility. Finally, we ran unbiased simulations of the complete apo and holo BRI1 and BRI1-BAK1 ECDs in order to examine the role of BRI1 LRRs 1-12 in the association process and in stabilizing the complex. In the holo complex, BRI1 underwent a large conformational change, forming a secondary interface with BAK1 through the BRI1 N-terminal LRRs.

## 2 Results

### BL binding stabilizes the tBRI1-BAK1 complex in a pH-dependent manner

Using a REUS sampling scheme, we calculated the standard free energy of tBRI1-BAK1 ECD association in the presence and absence of BL. We estimated the overall ΔG° of association values to be −2.120 ± 0.380 kcal·mol^−1^ for the apo complex and −3.360 ± 0.453 kcal·mol^−1^ for the holo complex (Table 1). The presence of BL in the island domain of tBRI1 stabilized the tBRI1-BAK1 complex by a moderate −1.510 ± 0.591 kcal·mol^−1^, and shifted the minimum in the separation PMF from ~26 Å to ~27 Å (Fig. 2). For both the apo and holo complexes, the separation PMFs were surprisingly shallow (Fig. 2). Individually, the free energies of adding and removing conformational restraints (Table 1) contributed far less than in previous studies [23], almost certainly due to the fact that we restrained RMSD values to non-zero values, giving tBRI1 and BAK1 some flexibility at the cost of longer convergence times.

**Table 1.**
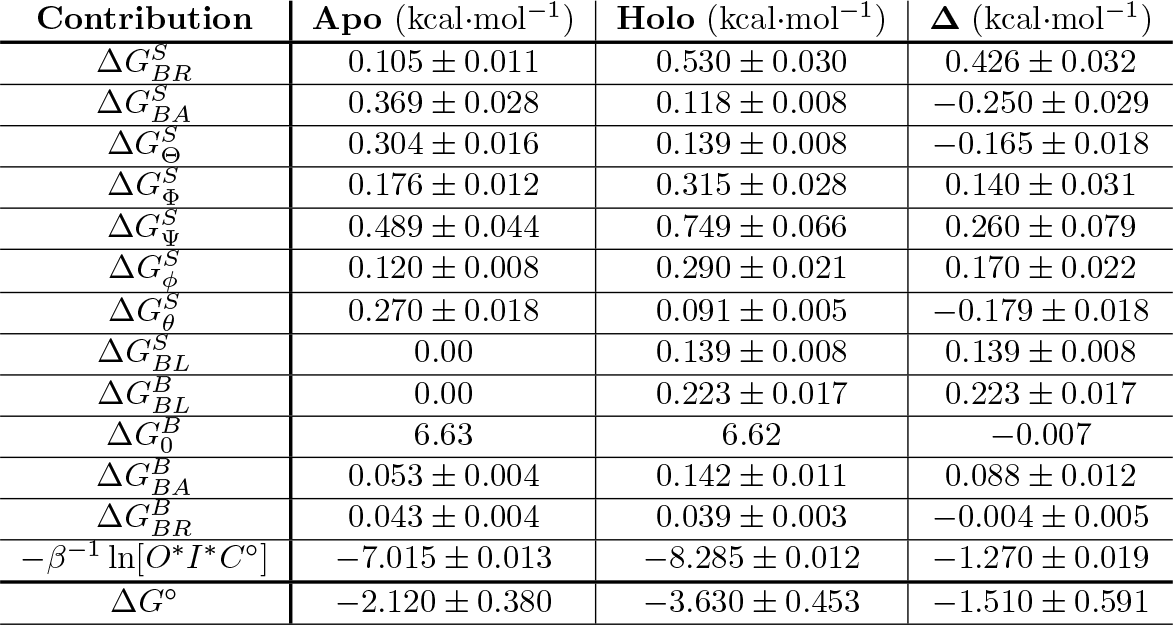
Calculated contributions to the free energy of BRI1-BAK1 association with standard deviations.

**Figure 1:**
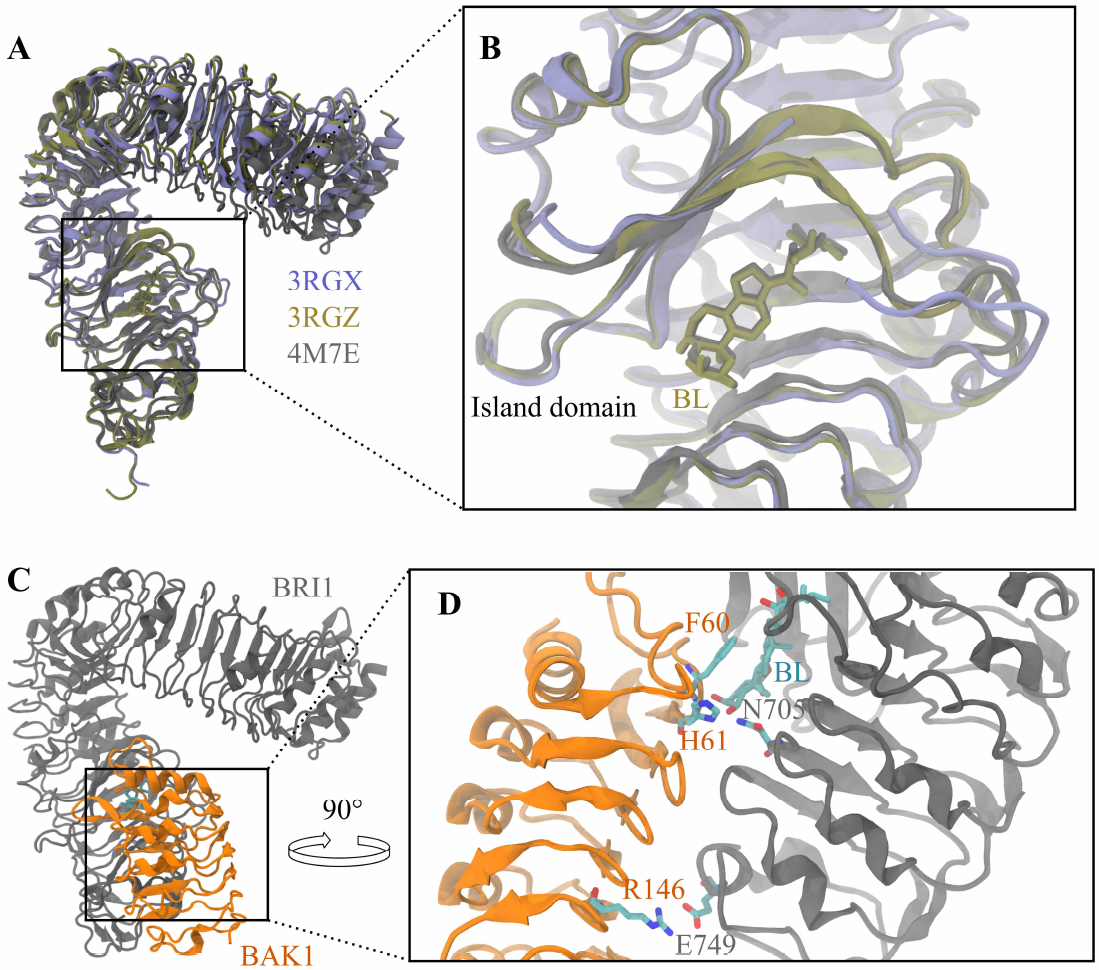
Analysis of BRI1 and BAK1 ECD crystal structures. A) Structural alignment of the BRI1 ECD from apo (PDBID: 3RGX [2], shown in purple), BL-bound (PDBID: 3RGZ [2], shown in tan), and BL-BAK1-bound (PDBID: 4M7E, chain A [7], shown in grey) crystal structures. B) A closeup of the island domains of structures shown in A). Note that part of the island domain of the apo structure (PDBID: 3RGX) is not resolved in the crystal structure, indicating disorder. C) Crystal structure of the BL-bound BRI1-BAK1 ECD complex (PDBID: 4M7E, chains A and C). BRI1 is shown in grey, while BAK1 is shown in orange. D) A closeup of the BRI1-BAK1-BL interface. The critical BAK1 residues F60 and H61 are shown interacting with BL and BRI1 N705. The BAK1 R146-BRI1 E749 interactions is also shown. This figure was produced using VMD 1.9.2 [59] and Inkscape 0.91 [60].

**Figure 2:**
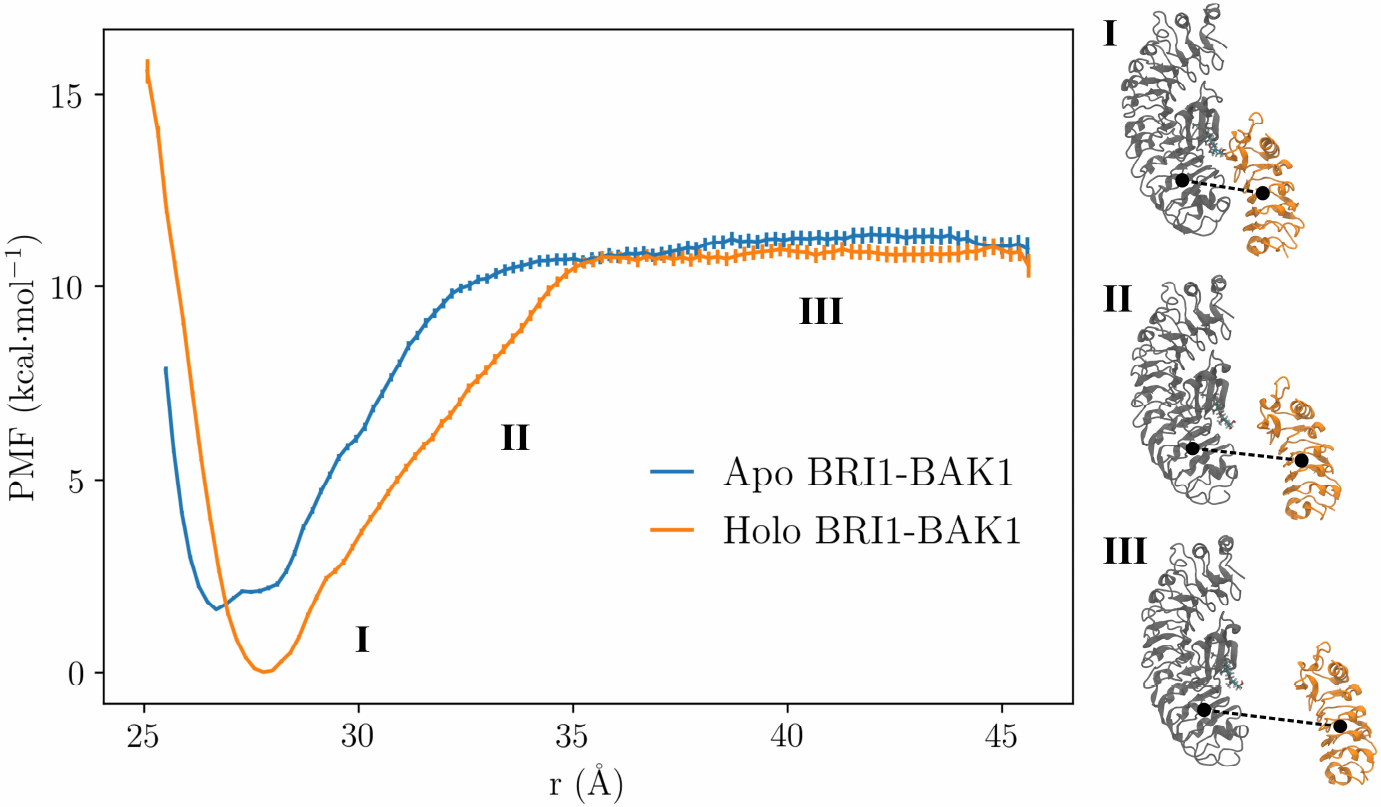
The potential of mean force as a function of BRI1-BAK1 separation distance (*r*) from REUS simulations. Roman numerals indicate associated (**I**), intermediate **II**), and dissociated (**III**) states. We estimated the PMF and error bars, representing standard deviations, using MBAR. This figure was produced using VMD 1.9.2 [59], Matplotlib 2.2.2 [61], and Inkscape 0.91 [60].

Results from analytical size exclusion chromatography have provided evidence that the ECDs of BRI1 and SERK1, a close homolog of BAK1, do not interact in the absence of BL [5]. This suggests a positive ΔG of association at relevant concentrations, in contrast to our ΔG° of −2.120 ± 0.380 kcal·mol^−1^. Previously reported experimental values of ΔG° for BAK1 association with BL-bound BRI1 are −8.44 kcal·mol^−1^ from grating-coupled interferometry and −9.20 kcal·mol^−1^ from isothermal titration calorimetry [19] (see Supplementary Information for details), both far larger in magnitude than our result of −3.630 ± 0.453 kcal·mol^−1^.

Both experimental association free energies were determined in solutions with pH 5 [19], whereas our protein models were constructed in the most likely protonation states at pH 7. BRI1 and BAK1 are known to interact more strongly in acidic conditions [6], possibly due to double protonation of the BAK1 H61 sidechain. In order to explore the origins of this pH dependence we performed alchemical free energy calculations in order to estimate ΔΔ*G* of association for apo and holo BRI1 and BAK1 with protonation of BAK1 H61 and the interfacial BRI1 E749.

For protonation of BAK1 H61 (Fig. S24 A-B), we estimated ΔΔ*G*_*Apo*_ = 2.729 ± 0.097 kcal·mol^−1^ and ΔΔ*G*_*Holo*_ = −1.958 ± 0.096 kcal·mol^−1^, the changes in tBRI1-BAK1 association free energy. Under the assumption that the internal and relative conformational dynamics of tBRI1 and BAK1 will not be significantly affected by BAK1 H61 protonation, we can add these values to our REUS Δ*G*° estimates, yielding 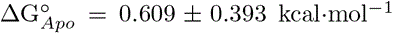 and 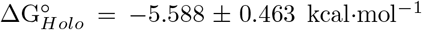 at pH 5. These estimates are far closer to the experimental values, reflecting the important role of BAK1 H61 protonation in distinguishing apo from holo BRI1.

### BL binding stabilizes the BRI1 island domain

It has been previously proposed that BR binding to the BRI1 ECD could alter the dynamics of the island domain, based on the decreased B-factors in the island domain of holo BRI1 ECD crystal structures as compared to apo BRI1 [2, 3, 20]. We estimated B-factors from conventional MD simulations of the apo and holo BRI1 ECDs according to 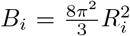[24], where *R*_*i*_ is the RMSF of residue *i*, and compared the *i* estimated B-factors to crystallographic B-factors [PDBIDs: 3RGX (apo) and 3RGZ (holo)] (Fig. 3). In the island domain region, the crystallographic B-factors are uniformly slightly higher for apo BRI1 than holo BRI1 (Fig. 3 A), as has been noted previously [2, 3, 20]. In comparison, B-factors calculated from our simulations of the BRI1 ECD did not display noticeable differences in the island domain between apo and holo systems (Fig. 3 B). The B-factors for both apo and holo BRI1 are much higher from our simulations than for the experimental systems, likely due to the fact that our simulations were performed at 300 K and in solution, in contrast to the conditions necessary for x-ray crystallography. Additionally, the particularly large B-factors in the terminal regions of BRI1 ECD from simulations suggest that there is likely large-scale flexibility in BRI1, which we further explore later.

**Figure 3:**
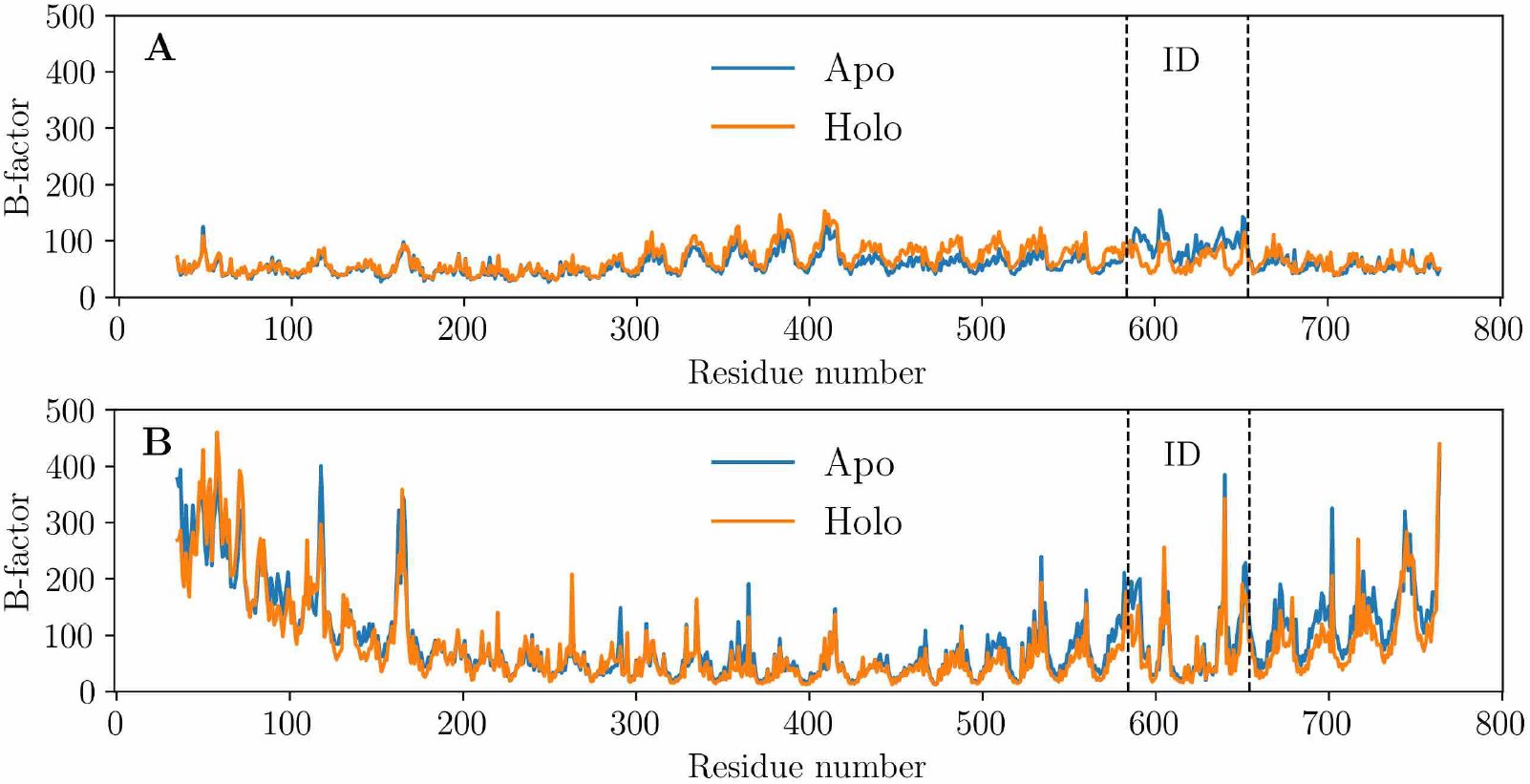
**A** Crystallographic B-factors for apo (PDBID: 3RGX [2]) and holo (PDBID: 3RGZ [2]) BRI1. The island domain is shown within the vertical dashed lines. Note the higher B-factors for the apo structure within the island domain. **B** B-factors calculated from unbiased molecular dynamics simulations of the apo and holo BRI1 ECD. The N-terminal and C-terminal regions display high B-factors as compared with their crystallographic counterparts, while there appears to be little difference between the apo and holo B-factors. This figure was produced using Matplotlib 2.2.2 [61].

As it is difficult to make conclusions from the B-factor calculations, we performed GAMD on apo and holo tBAK1 and calculated reweighted PMFs over the island domain RMSD with respect to a crystal structure (Fig. 4). Both the apo and holo tBRI1 island domain PMFs had a minimum near 1.2 Å, with a second local minimum at 1.7 Å (apo) and 1.9 Å (holo). Most notably, the apo PMF is far flatter than the holo PMF. At an RMSD of around 1.6 Å, the holo PMF is over 2 kcal·mol^−1^ larger than the apo PMF, and the difference in the PMFs largely increases at even larger RMSDs. These results indicate that binding of BL to the BRI1 island domain decreases fluctuations, stabilizing the island domain.

**Figure 4:**
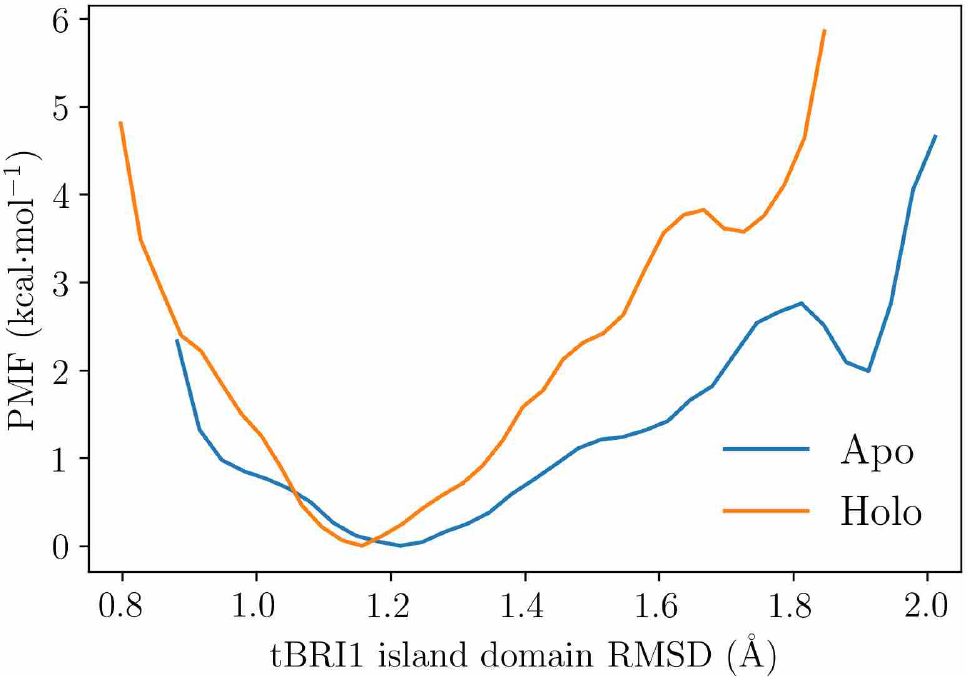
Potential of mean force for the RMSD of the apo and holo tBRI1 island domains with respect to a BRI1 ECD crystal structure (PDBID: 4M7E chain A [7]) from GAMD simulations. The holo island domain is stabilized with a single dominantly stable state at an RMSD of around 1.2 Å, while the apo island domain can attain a wider range of RMSDs. This figure was produced using Matplotlib 2.2.2 [61].

### BRI1 LRRs 1-12 are highly flexible and interact with BAK1

In order to examine the overall flexibility of the BRI1 ECD, we first performed anisotropic normal mode (ANM) analysis [25, 26] on the apo ECD of BRI1 using ProDy [27]. The structure of the apo BRI1 ECD was taken from the PDB 4M7E chain A, with BL removed. The first two modes, shown in Fig. 5, provide initial clues as to what global BRI1 dynamics look like. Mode I (Fig. 5 A-D) corresponds to relative motion between LRRs1-12 and the remainder of the ECD which takes BRI1 towards and away from a planar closed ring. Mode II (Fig. 5 E-H) corresponds to a curling motion, analogous to the coiling of a spring. The angle Ξ defined in the Methods section roughly corresponds to normal mode I while Ω roughly corresponds to normal mode II.

**Figure 5:**
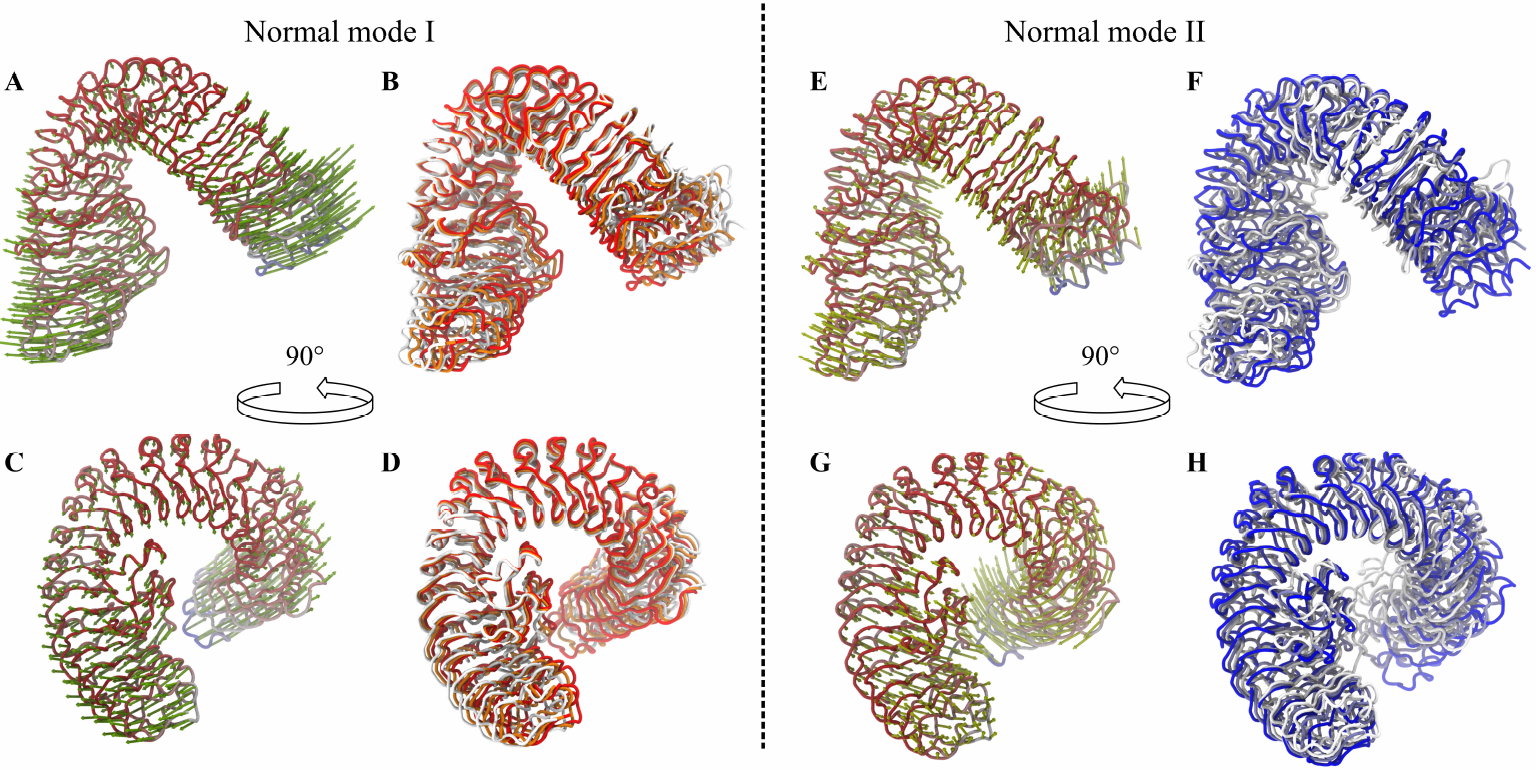
Anisotropic normal mode analysis of the BRI1 ECD taken from a crystal structure of the BRI1-BAK1-BL complex (PDBID: 4M7E chain A). **A-D** depict the first normal mode while **E-H** depict the second normal mode. **A** and **C** show the displacement vectors of *α*-carbons along normal mode I, where **C** is rotated 90° with respect to **A. B** and **D** depict conformations along the first normal mode, where again **D** is rotated 90° with respect to **B. E-H** can be described analogously to **A-D**. This figure was produced using VMD 1.9.2 [59], the VMD plugin Normal Mode Wizard [27], and Inkscape 0.91 [60]

Next, we performed unbiased MD simulations of the apo and holo BRI1 ECD and BRI1-BAK1 ECD complex, for a total of four systems. In addition to the angles Ξ and Ω described in the Methods section, we examined the distances between BRI1 R131 and BRI1 D651 for all four systems as well as the distance between BRI1 N108 and BAK1 K44 for the complexes. For the BRI1 ECD alone, LRRs 1-12 were flexible both in the presence and absence of BL. The BRI1 R131-D651 distance and both Ξ and Ω fluctuated around the same mean value in both apo and holo BRI1 in a manner qualitatively consistent with harmonic dynamics (Fig. 6), although the BRI1 R131-D651 distance distribution is apparently tighter for holo BRI1. While these results provide evidence that BRI1 LRRs 1-12 are in fact flexible, there appears to be little difference in apo and holo BRI1 ECD dynamics in the absence of the BAK1 ECD.

**Figure 6:**
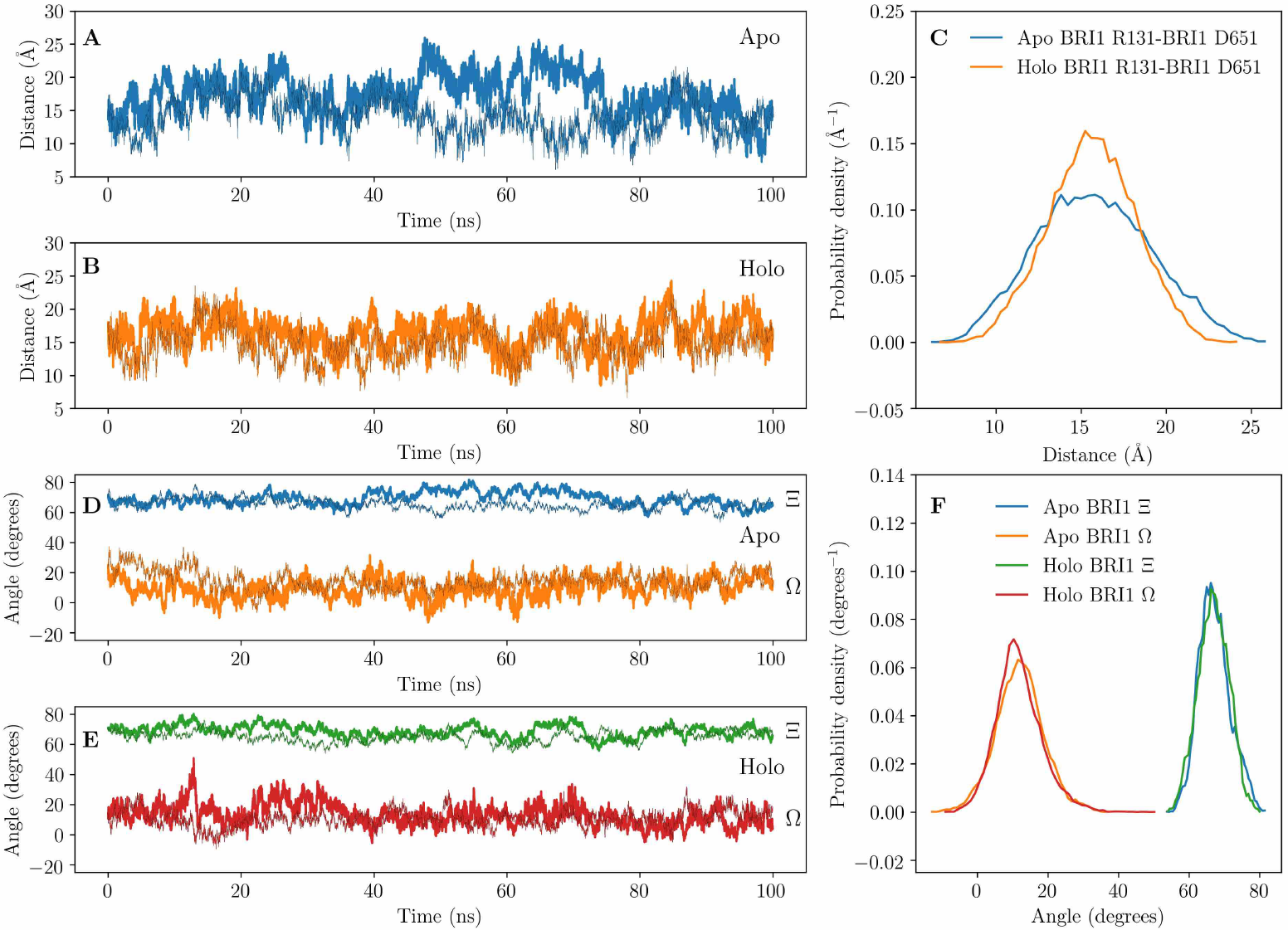
Conformational dynamics of the apo and holo BRI1 ECD. For all the following time series, two independent simulations are shown. **A** Time series of R131-D651 distance in apo BRI1. **B** Time series of R131-D651 distance in holo BRI1. **C** Probability densities of the R131-D651 distance in apo and holo BRI1 from the time series in **A** and **B** estimated with histograms. **D** Time series of Ξ and Ω in apo BRI1. **E** Time series of Ξ and Ω in holo BRI1. **F** Probability densities of Ξ and Ω in apo and holo BRI1 from the time series in **E** and **F** estimated with histograms. This figure was produced using Matplotlib 2.2.2 [61].

In simulations of the BRI1-BAK1 ECD complex, we found that without BL bound, the dynamics of BRI1 differed very little from BRI1 dynamics without BAK1 present (Fig. 7). However, in the presence of BL, BRI1 underwent a conformational change after around 26 ns of simulation, forming a closed BRI1 conformation featuring a second BRI1-BAK1 interface. The closed conformation is characterized by formation of a contact within BRI1 between R131 and D651, which remained stable for the remaining 34 ns of the trajectory after the conformational change occurred (Fig. 7). A contact between BRI1 N108 to BAK1 K44 also formed, although this interaction was able to break and reform several times (Fig. 7 C) even after the conformational change, leading to a multi-peaked distribution (Fig. 7 D). This conformational change corresponds with a change in BRI1 Ξ (Fig. 7 C-D), demonstrated by the stability of Ξ at around 40° starting around the 26 ns mark and by the doubly peaked angle distribution. This suggests that this conformational change occurs roughly along normal mode I, while motion along normal mode II does not contribute as much, although Ω appears to be stabilized in the holo complex especially after the conformational change (Fig. 7 C-D).

**Figure 7:**
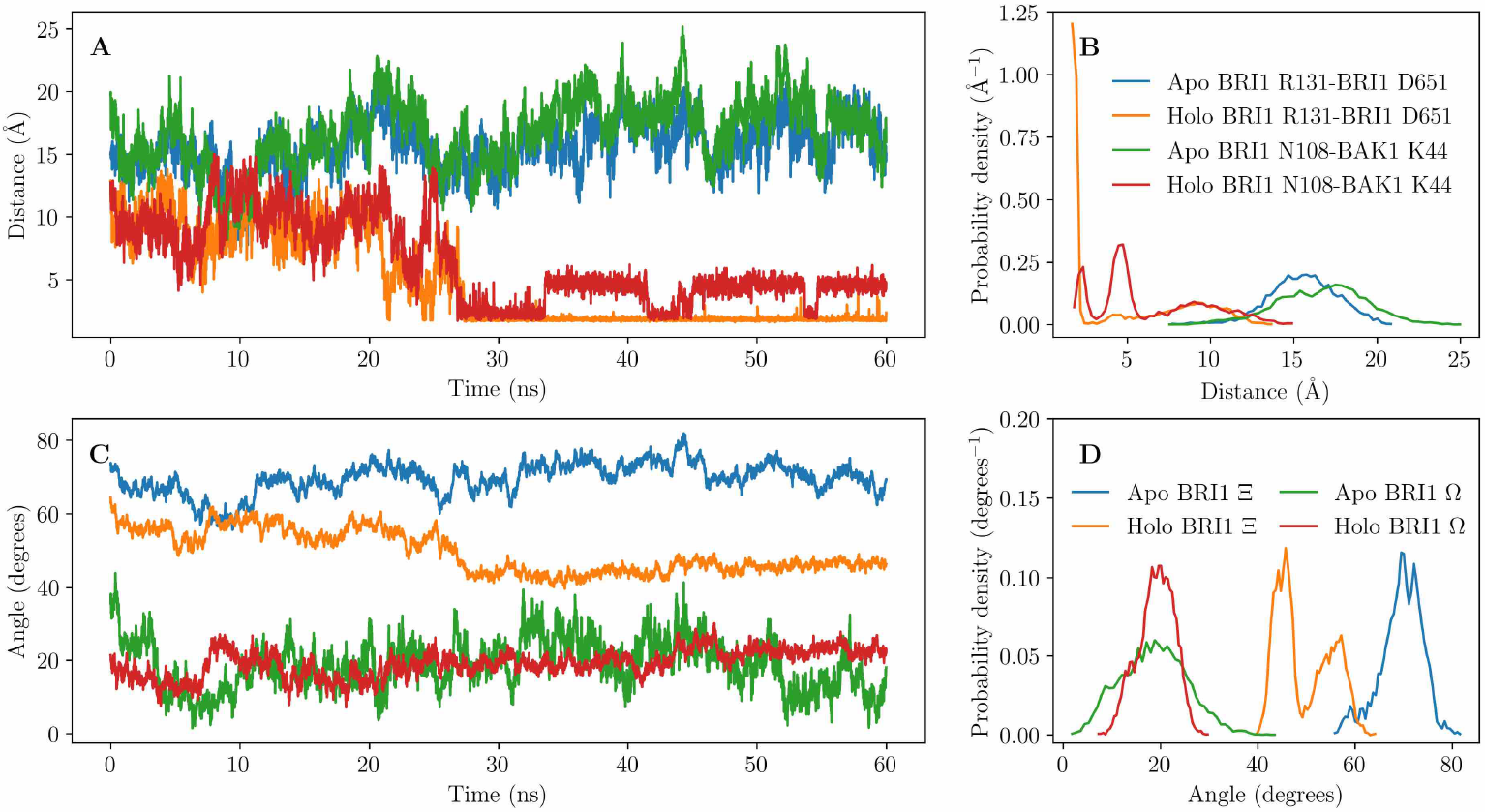
Conformational dynamics of the apo and holo BRI1-BAK1 ECD complex. **A** Time series of the BRI1 R131-D651 and BRI1 N108-BAK1 K44 distances in apo and holo systems. Note the conformational change near 26 ns in the holo system, and the subsequent fluctuations between two apparent states in the BRI1 R131-D651 and BRI1 N108-BAK1 K44 distance. **B** Probability densities of the BRI1 R131-D651 and BRI1 N108-BAK1 K44 distances in apo and holo systems estimated using histograms. **C** Time series of Ξ and Ω in the apo and holo BRI1-BAK1 complexes. **D** Probability densities of Ξ and Ω in the apo and holo BRI1-BAK1 complexes estimated using histograms. This figure was produced using Matplotlib 2.2.2 [61].

As mentioned before, in the secondary interface BRI1 D651 interacts with BRI1 R131 and BAK1 K44 interacts with BRI1 N108 as well as several backbone oxygens (Fig. 8). Additionally, BRI1 N37 interacts with the backbone oxygen of BRI1 S184 (Fig. 8). We predicted the mean and mode of the BRI1 D651 pKa distribution from BRI1 apo simulations to be 3.712 and 3.944 respectively, using PROPKA 3.1 [28, 29], meaning that this interaction is likely as strong at pH 5 as it is at pH 7.

**Figure 8:**
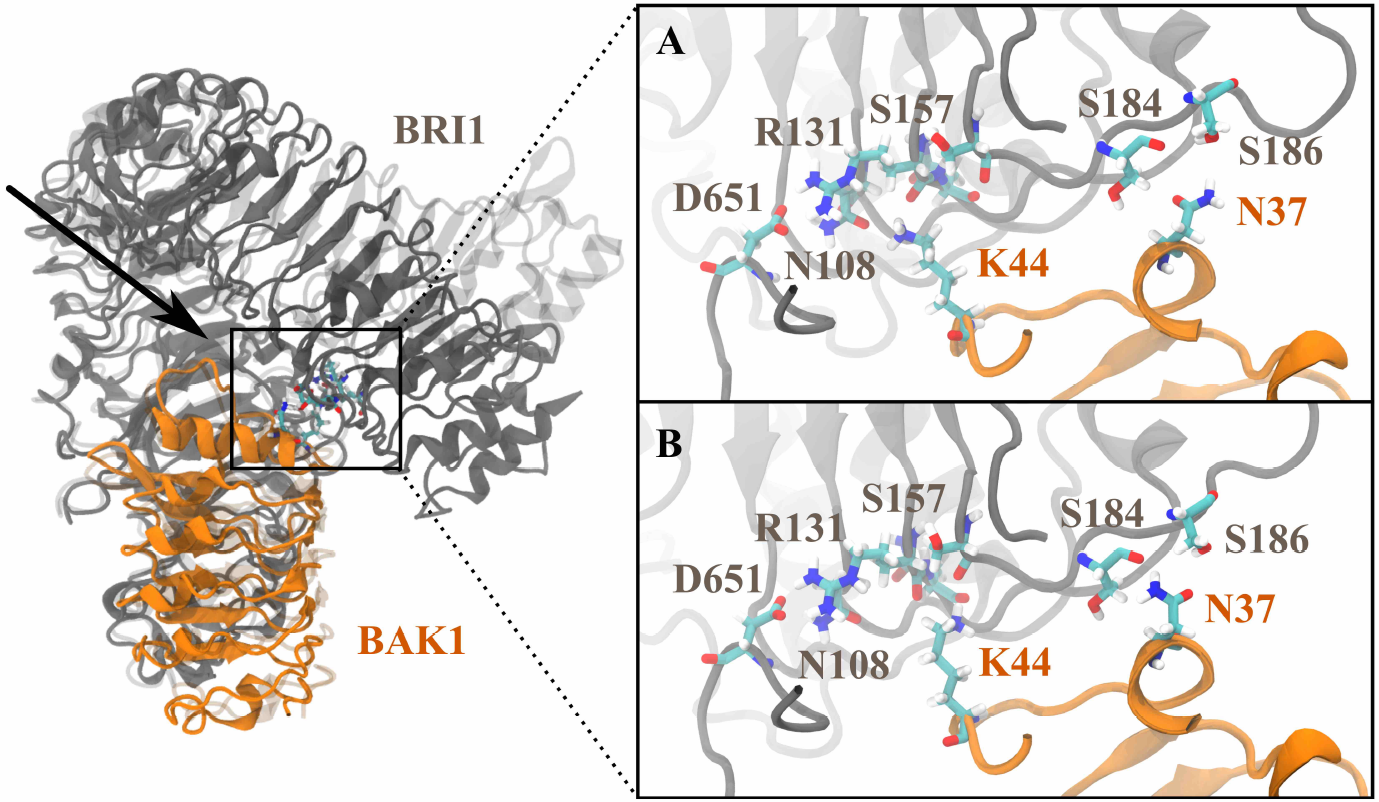
The secondary interface between the BRI1 and BAK1 ECDs. The crystal structure (PDBID: 4M7E chains A and C [7]) is shown in a transparent representation. The inset subfigures **A** and **B** represent two alternative interfaces see in simulations, with **A** being the state in Fig. 7 C with BRI1 N108 and BAK1 K44 at the smallest distance and **B** being the state with a larger BRI1 N108-BAK1 K44 distance, where BAK1 K44 interacts with BRI1 backbone oxygens.

It is important to note that we cannot make claims with any confidence regarding the thermodynamics of BRI1 conformational changes from such short simulations. We have found that the closed BRI1 conformation can occur in simulations of the holo BRI1-BAK1 complex within a timespan of 60 ns, but it is difficult to make any statement about the stability of this conformation from our data. Similarly, BRI1 LRRs 1-12 are flexible, but the presented histograms in Fig 6 likely only represent local BRI1 dynamics and may not represent the global dynamics of the BRI1 ECD. Further investigation of BRI1 ECD dynamics is required, but is beyond the scope of this study.

## 3 Discussion

Our examination of three mechanisms of BL-induced BRI1-BAK1 association indicates that each one likely plays an important role. Our REUS-based Δ*G*° estimates indicate that at pH 7 the BRI1-BAK1 ECD complex is weakly favored both with and without BL bound. Alchemical free energy calculations suggest that protonation of BAK1 H61, consistent with a solution pH of 5, is instrumental in differentiating apo and holo BRI1, increasing the stability gap between the holo and apo BRI1-BAK1 complex from −1.510 ± 0.591 kcal·mol^−1^ to −6.197 ± 0.607 kcal·mol^−1^. With the contributions of BAK1 H61 protonation, we estimate tBRI1-BAK1 ECD association free energies of 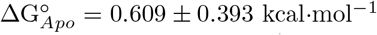 and 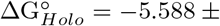 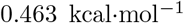, closer to the experimental values of −8.44 kcal·mol^−1^ or −9.20 kcal·mol^−1^ for the holo complex, but still less stable by several kcal·mol^−1^. Our GAMD simulations provide direct evidence that BL binding to BRI1 stabilizes the island domain, likely reducing the entropic cost of BRI1-BAK1 association. Finally, unbiased MD simulations revealed a conformational change in the BRI1 ECD when complexed with BAK1 and BL, forming a secondary interface between BRI1 and BAK1, which may further stabilize the BRI1-BAK1 complex.

Are our results consistent with experimental findings? The combined contributions of direct BL-BAK1 interactions, BL-induced BRI1 island domain stabilization, and formation of the second BRI1-BAK1 interface complicate direct comparison of our REUS-based association free energies with experimental quantities. Beyond the inaccuracies of forcefields in describing protein-protein interaction [30], and the possibility of insufficient sampling, which does not appear to be an issue (Fig. S1), we found that direct BL-BAK1 interactions contribute significantly to the overall BRI1-BAK1 BL-dependent association free energy, but cannot entirely account for BRI1-BAK1 association. From alchemical free energy calculations, protonation of BAK1 H61 appears to play a critical role in distinguishing apo and holo BRI1, destabilizing the apo BRI1-BAK1 complex by −2.729 ± 0.096 kcal·mol^−1^ while stabilizing the holo complex by 1.958 ± 0.096 kcal·mol^−1^. The important role of pH in the stability of the BRI1-BAK1 complex is consistent with experimental findings [6].

Although it appears direct interactions between BRI1 and BAK1 mediated by BL, that is, the molecular glue hypothesis, does not account for the entire association free energy, it is difficult to claim with certainty that the difference between our estimates and the experimental association free energies implies other sources of BRI1-BAK1-BL complex stability. The fact that we observe stabilization of the BRI1 island domain upon BL binding and formation of a second BRI1-BAK1 interface with BL bound adds to the weight of evidence suggesting that the molecular glue hypothesis alone cannot explain BL-induced BRI1-BAK1 association. Although we did not observe a conformational change in apo BRI1 complexed with BAK1, we cannot be sure that BRI1 is less likely to undergo conformational change without BL bound than it is with BL bound. However, as BAK1 is far less likely to associate with apo BRI1 than holo BRI1, even an equally probable BRI1 conformational change in the apo and holo complexes would preferentially stabilize the favorable holo complex (Fig. 9).

**Figure 9:**
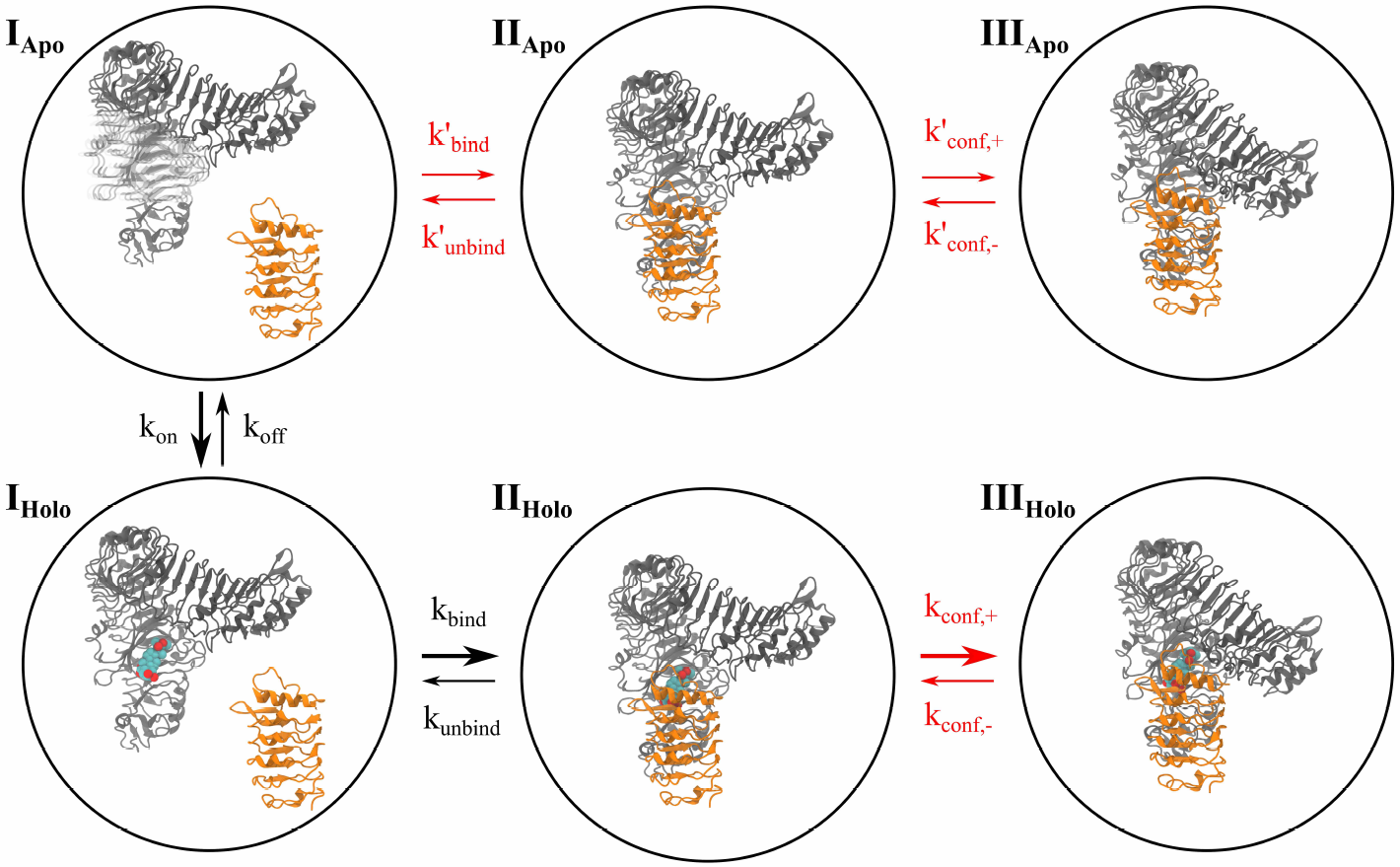
A model of BRI1-BAK1 BL-induced ECD association. Rates in black have been experimentally determined, while rates in red have not. The thickness of each arrow represents the magnitude of corresponding rate. Through the arrow thicknesses, we imply that 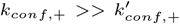, although this claim is only weakly supported by our simulations and requires further investigation. This figure was produced using VMD 1.9.2 [59] and Inkscape 0.91 [60].

Experimental evidence already exists for the molecular glue and island domain stabilization hypotheses. In this paper, we have made the additional claim that BRI1 LRRs 1-12 are highly dynamic and play an important role in stabilizing the BRI1-BAK1 ECD complex. Supposing that this claim is true, BRI1 mutations D651A and R131A should increase the effective experimental BRI1-BAK1 dissociation rate, *k*_*d*_, and consequently decrease *K*_*A*_. This should occur because BRI1 R131 and D651 interact and stabilize the closed BRI1 conformation and mutating either will destabilize the closed BRI1 conformation. Furthermore, BAK1 mutations N37A and K44A or BRI1 mutations N108A or D161A will increase BRI1-BAK1 *k*_*d*_, while the double mutations BAK1 N37A-K44A or BRI1 N108A-D161A will further increase *k*_*d*_. This should occur because any of these mutations will disrupt interactions in the secondary BRI1-BAK1 interface, which is only formed when BRI1 is in the closed conformation.

LRR-RLKs are an important group of receptors in plant cells with a common evolutionary ancestry [31] and the shared structural features [32, 20], namely the predominance of LRR domains in the ECD. Our finding that BRI1 undergoes a conformational change in order to form a second interface with BAK1 raises several questions about the LRR-RLKs in general. How flexible are the ECD domains of larger LRR-RLKs? To what extent does this flexibility play a functional role in association with a coreceptor and receptor activation? How common is the formation of secondary interfaces between larger BRI1-like and smaller BAK1-like LRR-RLKs? We believe that a combination of computational modeling and biophysical experiments is uniquely suited to address these questions, which could have significant impact on our understanding of plant growth, developmental, and immune signaling.

In summary, we find that direct BL-BAK1 interactions (the molecular glue hypothesis), BRI1 island domain stabilization, and a secondary BRI1-BAK1 interface all likely play a role in BRI1-BAK1 association. BL-induced association of the BRI1 and BAK1 ECDs may occur through the following sequence of events (Fig. 9): BL binds to the BRI1 island domain, stabilizing it in a conformation amenable to BAK1 association. With the BRI1 LRRs 1-12 in a crystal structure-like conformation, BAK1 associates with BRI1 and interacts with BL through F60, H61, V62, and D74. BRI1 LRRs 1-12 can then undergo a conformational change, forming the secondary BRI1-BAK1 interface and further stabilizing the BRI1-BAK1 complex.

## 4 Methods

### System setup

All systems were set up using the tleap program within AmberTools 15 [33] using the Amber ff14SB force field [34]. Initially, the protein atoms of the BRI1-BAK1 extracellular domains in complex with BL were taken from a crystal structure (PDBID: 4M7E, chains A and C [7]). Histidine protonation states were determined in accordance with a solution at pH 7 using the H++ 3.2 webserver [35]. Disulfide bonds were added according to the crystal structure. All systems were solvated in boxes of TIP3P [36] water molecules, so that the edges of the box were at least 10 Å from any protein atom. Parameters for BL were assigned using the Antechamber program within AmberTools15 and the GAFF forcefield[37]. As Antechamber assigns a small, non-integer overall negative charge to BL, the absolute value of the overall charge was distributed as evenly as was practical across all BL atoms so as to obtain an overall neutral charge. Sodium and chloride ions were added to each system so as to neutralize the charge and bring the salt concentration to ~150 mmolal NaCl. Simulations using the full BRI1 ECD consisted of residues 34-766, while the tBRI1 ECD consisted of residues 378-766, and the BAK1 ECD consisted of residues 26-200.

### General simulation details

All simulations, except alchemical free energy calculations (see below), were run in NAMD 2.12-2.13 [38] with a time step of 2 fs and with hydrogen-containing bonds constrained using the SHAKE algorithm [39]. All production runs were maintained at constant temperature of 300 K using a Langevin thermostat with a coupling time constant of 2 ps and at a constant pressure of 1 bar using a Berendsen barostat. The particle mesh Ewald method [40] was used to treat electrostatics and a cutoff of 10 Å was used for non-bonded interactions. Trajectory analysis was performed using MDTraj [41] and NumPy [42] within Jupyter Notebooks [43].

### Determining BRI1-BAK1 extracellular domain *K*_*A*_s

We calculated the equilibrium association constant (*K_A_*) of BAK1 to BRI1 with and without BL bound to BRI1 using the replica-exchange umbrella sampling (REUS)[44, 45] approach developed by Woo and Roux[46] and Gumbart *et al.* [47, 23]. For the apo complex, five restraints on the relative orientation of the BRI1 and BAK1 ectodomains were introduced in addition to the restraint on the PMF CV (Figure 10). An additional restraint was added in the holo complex to ensure that BL remained bound to BRI1 (Figure 10).

**Figure 10:**
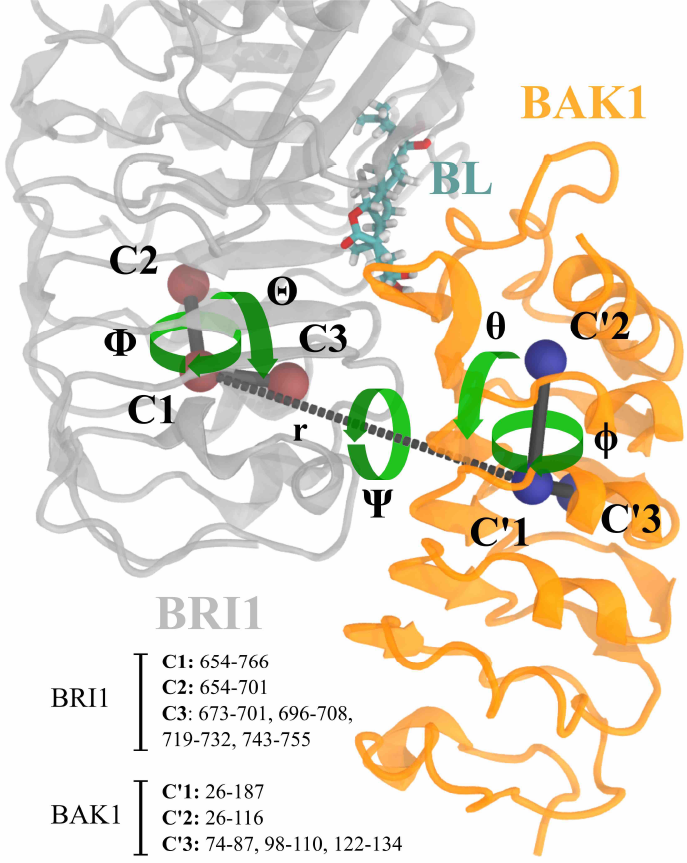
Collective variables harmonically restrained during umbrella sampling. C1 to C3 and C’1 to C’3 are the centers of mass of the *α*-carbons of different sets of residues in BRI1 and BAK1, respectively (the residues used are listed in the lower lefthand corner). The distance *r* between C1 and C’1 was used as the CV for umbrella sampling. The angular restraints are defined as: Θ (C’1-C1-C2) and *θ* (C1-C’1-C’2). The torsion restraints are defined as: Φ (C’1-C1-C2-C3), *ϕ* (C1-C’1-C’2-C’3), and ψ (C2-C1-C’1-C’2). Additionally, for the BRI1-BL-BAK1 simulations, the distance between the center of mass of all heavy atoms in BL and the *α*-carbon of BRI1 F658 was restrained to prevent BL unbinding. This figure was produced using VMD 1.9.2 [59] and Inkscape 0.91 [60].

#### Replica-exchange umbrella sampling

Both the apo and holo BRI1-BAK1 complexes were set up as described above, subjected to 20,000 steps of energy minimization and equilibrated for 2 ns. The final 1 ns of equilibration for both systems was used to compute average values of each restraint CV in the complex. These average values were used as the reference values of each CV in the corresponding harmonic restraints (Supplementary Table [REF]). Next, the center of the C1-C’1 distance restraint was increased from the equilibrium value of both associated complexes to 45 Å over 1 ns simulations in order to generate starting structures for each umbrella sampling window. REUS along the C1-C’1 distance was performed with a total of 61 windows evenly spaced between 25 Å and 45 Å (0.33 Å intervals between window centers), where the starting structure for each window was equilibrated for 1 ns and each replica was simulated for 7 ns. An additional 7 ns of REUS was performed for each window of both the apo and holo systems to ensure convergence of the separation PMF (Fig. S1). A total of 854 ns of REUS was performed in order to estimate the separation PMF of each system.

We used REUS to calculate the free energies of applying and removing restraints on the relative orientations of BRI1 and BAK1 (Θ, Φ, *θ*, *ϕ*, and ψ) in addition to RMSD and BL position restraints in both the apo and holo states. In order to generate wide ranges of each restrained CV for initiating each umbrella sampling window, we performed temperature-accelerated MD[48, 49], coupling each respective CV to a dummy particle experiencing a temperature of 2500 K. The number of windows for the REUS simulations of each CV varied, while each window was equilibrated for 1 ns, followed by 7 ns of REUS.

Rather than focusing on the PMFs estimated from each set of REUS simulations, we examined the integrands of the target ensemble averages for convergence (Fig. S4-S23). We ensured that each integrand qualitatively converged with time and that the integrand fell to zero at both edges of the sampled range for each restraint CV, indicating that the ensemble averages should be estimated well from our simulations. For the RMSD restraint on apo BAK1 in the bound state with respect to BRI1, we noticed that the range of RMSD sampled was insufficient to estimate the ensemble average, and subsequently performed steered MD followed by standard umbrella sampling for an additional 6 windows (Fig. S5).

#### PMF estimation

All unbiased PMFs were calculated using the multistate Bennett acceptance ratio (MBAR) estimator [50] as implemented in pymbar 3.0.4 [51]. All trajectories were subsampled to a frequency of 50 ns^−1^ to reduce correlations between samples while maintaining a sufficient number of samples (Fig. S2-S3). While this choice should improve the PMF estimates over samples chosen according to the correlation time [52] due to retention of more samples, it likely also resulted in underestimation of the standard deviations for bin free energies due to sample correlation [50].

#### Overall *K*_*A*_ calculations

As per [47, 23], the overall standard dissociation constant can be calculated as a product of terms, each calculated from simulations or analytically in the case of bulk alignment contributions. For simulations in the bound state (referred to as “site” in [47, 23]), the distance between C1 and C’1 (Figure 10), *r*, was subjected to a harmonic restraint with a force constant of 10 kcal·mol^−1^Å^−2^ if *r* increased beyond 29 Å. For simulations in the bulk state, a distance of *r*\* = 44 Å was chosen according to a distance where both separation PMFs had plateaued, and *r* was harmonically restrained at *r*\* with a force constant of 10 kcal·mol^−1^Å^−2^.

The overall calculation of Δ*G*° of association involves calculating the free energy of adding each restraint one after another in the bulk state, calculating the free energy of translating the restrained BRI1 and BAK1 to the bound state, and then calculating the free energy of releasing each restraint in the bound state. The free energies of adding and removing restraints can conveniently be calculated using averages of exponentials of restraint potential energy (see Supplementary Information for full details).

The separation PMF integrals were approximated as

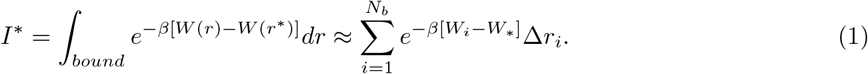

where *W* (*r*) is the PMF at separation distance *r*, *N*_*b*_ is the number of bins used to approximate *W* (*r*) in the bound state, and *W*_*i*_ is the free energy of occupying bin *i*. The bound region was defined as the set of 13 bins centered on the bin with the lowest overall PMF (roughly *r*_0_ ± 1.32 Å for the apo system and *r*_0_ ± 1.35 Å for the holo system). Assuming mutual independence of each *W*_*i*_, we approximated the variance of *I*\* 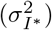 using a Taylor series expansion truncated at the first order term as

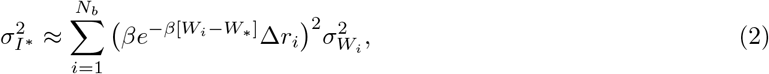

where 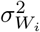 is the MBAR variance of *W*_*i*_. The exponential averages needed to calculate restraint free energy contributions were estimated as MBAR expectation values.

### BAK1 H61 protonation

We performed alchemical free energy perturbation simulations on snapshots from separation PMF REUS windows 8 and 57 (corresponding to separation distances of 27.67 Å and 44 Å, respectively) in order to assess the contributions of residue protonation to association free energy. We focused on the protonation of BAK1 H61, which interacts with BRI1-bound BL and has been proposed to be an important factor in pH-dependent BRI1-BAK1 interaction [6]. Because BRI1 E749 is in the BAK1-BRI1 ECD interface and initially appeared to have a pK_a_ above 5 based on predictions from H++ [35] on a crystal structure (PDBID: 4M7E), we also estimated the free energy of BRI1 E749 protonation through alchemical free energy calculations. However, PROPKA 3.1 [28, 29] calculations based on the whole apo BRI1 ECD simulations suggest that BRI E749 would probably be unprotonated at pH 5 (Fig. S25), meaning that BRI1 E749 protonation analysis is not particularly pertinent.

All alchemical free energy simulations were performed using Amber 18 [53, 54], using a Monte Carlo barostat. Due to the change in charge that occurs when protonating a residue, we included a co-alchemical Na^+^ ion in each simulation to maintain a net zero system charge [55], where we linearly scaled the charge to zero concurrently with linearly scaling the amino acid proton charge to one. No restraints on the co-alchemical ions were used. We used 41 windows in each alchemical transformation, where each window was run for 11 ns with the first 1 ns discarded as equilibration. In order to prevent association of BRI1 and BAK1, we applied restraints on the *α*-carbons of BRI1 residues 651-666 and of BAK1 residues 86-90 in all simulations started from snapshots with *r* ≈ 44 Å. The free energy of protonation for each case was estimated using MBAR, with frames sampled at a rate of 100 ns^−1^.

### Accelerated MD simulations of apo and holo tBRI1

We performed dual boost Gaussian accelerated MD (GAMD) [56] simulations of the apo and holo BRI1 ECD using NAMD 2.13 [57]. Restraints were added to the terminal regions of tBRI1 to prevent unrealistic unfolding and on the overall orientation of tBRI1 to allow for the use of a smaller periodic box. Each system was subjected to energy minimization for 20,000 steps, followed by 6 ns of equilibration. In the initial phase of simulations, potential energy statistics were recorded during an additional 11 ns of simulation, after which GAMD biases were applied. Each system was equilibrated for 1 ns with GAMD biases on according to the gathered statistics, after which 11 ns of further equilibration was performed where the bias parameters *E* and *k*_0_ were updated. In the final phase, ten independent GAMD trajectories were run for both apo and holo BRI1 each for a total of ~90 ns, saving frames every 200 fs. Each trajectory was run in six 15 ns segments, where the energy statistics and bias parameters were updated in the first and second nanoseconds of simulation, respectively, each with 400 ps of preparation simulations. While possibly not ideal for cumulant expansion reweighting, the entire 90 ns of each trajectory was used for final analysis. Unbiased PMFs were estimated using second-order cumulant expansions [58]:

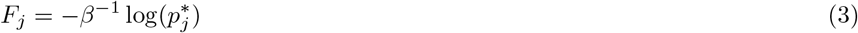

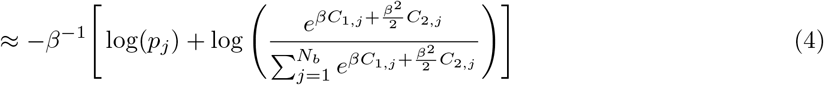

where *F*_*j*_ is the reweighted free energy of occupying bin *j*, *p*_*j*_ is the unreweighted histogram count of bin *j*, and *C*_*i,j*_ is the *i*th cumulant of the GAMD boost energy for frames in bin *j*. Bins with counts fewer than 100 were removed from the final PMFs.

### BRI1 conformational analysis

Two angles, Ξ and Ω, were defined in order to explore the conformational dynamics of the entire BRI1 ECD. We defined four centers of mass, from the backbone atoms of residues 34-104 (**c**_1_ ∈ ℝ^3^), 253-293 (**c**_2_), 383-423 (**c**_3_), and 667-767 (**c**_4_). We then defined **c**_21_ as **c**_2_ − **c**_1_, **c**_32_ as **c**_3_ − **c**_2_, and **c**_34_ as **c**_3_ − **c**_4_. The angle Ξ was defined as

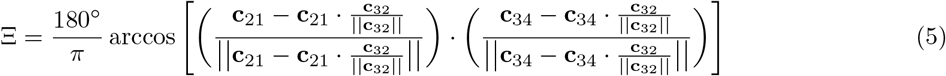

and the angle Ω as

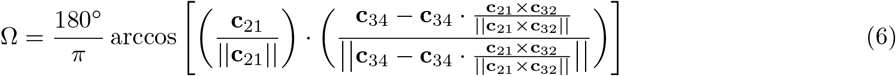

where caution was taken to differentiate distinct angles yielding identical values of Ξ and Ω, allowing for negative values of each angle.

We ran unbiased simulations of the apo and holo complete BRI1 ECD, as well as the apo and holo BRI1-BAK1 ECD complex. Two 100 ns simulations were run for each of the apo and holo BRI1 systems, while a single 60 ns simulation was run of both the apo and holo BRI1-BAK1 complex.

## Supporting information

Supplementary Methods, Images, Tables and Results

## 5 Acknowledgements

We thank the Blue Waters sustained-petascale computing project for providing computing time for this study. The Blue Waters sustained-petascale computing project is supported by the National Science Foundation (awards OCI-0725070 and ACI-1238993) and the state of Illinois.

## 7 Author Contribution

A.S.M. and D.S. designed the research. A.S.M. performed the research and analyzed data. A.S.M. wrote the manuscript with input from D.S.

