## Supplementary Methods, Images, Tables and Results for "How do brassinosteroids activate their receptors?"

### BRI1 E749 protonation

For the sake of completeness, we report results for BRI1 E749, although BRI1 E749 is unlikely to be protonated at pH 5 (Fig. S25). Counterintuitively, protonation of tBRI1 E749 appeared to strongly stabilize both the apo and holo tBRI1-BAK1 complex (Fig. S24 C-D). As BRI1 E749 interacts with BAK1 R146 in crystal structures (Fig. 1), we expected protonation of BRI1 E749 to break this interaction and weaken the stability of the rBRI1-BAK1 complex. However, for protonation of tBRI1 E749 we estimated $\Delta\Delta G_{Apo} = -12.225 \pm 0.101$ kcal·mol$^{-1}$ and $\Delta\Delta G_{Holo} = -7.113 \pm 0.102$ kcal·mol$^{-1}$. It appears that the interaction partners of BRI1 E749 in BAK1 are involved in unrealistic interactions with the truncated C-terminus of tBRI1.

### Calculation of association free energies from previously reported dissociation constants

Here, we show our calculations for binding free energy given the dissociation constants obtained by Hohmann and coworkers [1]. We assume a temperature of 300 $K$ and use a standard concentration of $C^\circ = 1/(1661\ \text{Å}^3)$. The standard free energy of association between the BAK1 and BL-bound BRI1 extracellular domains was calculated from grating-coupled interferometry results as follows:

$$K_D = 0.71\ \mu M = 0.71 \frac{\mu\text{mol}}{L} \cdot \frac{1\ \text{mol}}{10^6\ \mu\text{mol}} \cdot \frac{1\ L}{10^{27}\ \text{Å}^3} \cdot 6.022 \cdot 10^{23}\ \text{mol}^{-1} \approx 4.28 \cdot 10^{-10} \text{Å}^{-3}$$

$$K_A = \frac{1}{K_D} \approx 2.34 \cdot 10^9\ \text{Å}^3$$

$$\Delta G^\circ = -RT \log(K_A / 1661\ \text{Å}^3) \approx -8.44\ \text{kcal} \cdot \text{mol}^{-1}$$

The free energy of association from isothermal titration calorimetry (-9.20 kcal·mol$^{-1}$) was calculated accordingly.

### Calculation of association free energies from REUS

For the sake of brevity, we use the following shorthand notation for averages: $\langle A(\mathbf{x}) \rangle^E_{\{\alpha,\beta,\gamma\dots\}}$, where $A(\mathbf{x})$ is the quantity, dependent on protein coordinates, to be averaged, the superscript $E$ denotes the state of the association progress, either at the binding site ($S$) or in bulk solution ($B$), and $\{\alpha, \beta, \gamma \dots\}$ represents the set of collective variables restrained when the average is taken. For example,

$$\langle e^{-\beta u_\Theta(\mathbf{x})} \rangle^S_{\{BR,BA\}} = \frac{\int_S e^{-\beta[U(\mathbf{x})+pV(\mathbf{x})+u_{BR}(\mathbf{x})+u_{BA}(\mathbf{x})+u_\Theta(\mathbf{x})]} d\mathbf{x}}{\int_S e^{-\beta[U(\mathbf{x})+pV(\mathbf{x})+u_{BR}(\mathbf{x})+u_{BA}(\mathbf{x})]} d\mathbf{x}} \tag{1}$$

is the average value of $e^{-\beta u_\Theta(\mathbf{x})}$ in the $NPT$ ensemble, $u_\Theta(\mathbf{x})$ being the restraint potential on $\Theta$, with BRI1 and BAK1 bound and with the BRI1 and BAK1 backbone atoms restrained. The overall calculations of standard association free energies proceed as follows:

$$\Delta G^S_{BR} = \beta^{-1} \ln \left[ \langle e^{-\beta u_{BR}(\mathbf{x})} \rangle^S \right] \tag{2}$$
$$\Delta G^S_{BA} = \beta^{-1} \ln \left[ \langle e^{-\beta u_{BA}(\mathbf{x})} \rangle^S_{\{BR\}} \right] \tag{3}$$
$$\Delta G^S_{\Theta} = \beta^{-1} \ln \left[ \langle e^{-\beta u_{\Theta}(\mathbf{x})} \rangle^S_{\{BR,BA\}} \right] \tag{4}$$
$$\Delta G^S_{\Phi} = \beta^{-1} \ln \left[ \langle e^{-\beta u_{\Phi}(\mathbf{x})} \rangle^S_{\{BR,BA,\Theta\}} \right] \tag{5}$$
$$\Delta G^S_{\Psi} = \beta^{-1} \ln \left[ \langle e^{-\beta u_{\Psi}(\mathbf{x})} \rangle^S_{\{BR,BA,\Theta,\Phi\}} \right] \tag{6}$$
$$\Delta G^S_{\phi} = \beta^{-1} \ln \left[ \langle e^{-\beta u_{\phi}(\mathbf{x})} \rangle^S_{\{BR,BA,\Theta,\Phi,\Psi\}} \right] \tag{7}$$
$$\Delta G^S_{\theta} = \beta^{-1} \ln \left[ \langle e^{-\beta u_{\theta}(\mathbf{x})} \rangle^S_{\{BR,BA,\Theta,\Phi,\Psi,\phi\}} \right] \tag{8}$$
$$\Delta G^S_{BL} = \beta^{-1} \ln \left[ \langle e^{-\beta u_{BL}(\mathbf{x})} \rangle^S_{\{BR,BA,\Theta,\Phi,\Psi,\phi,\theta\}} \right] \tag{9}$$
$$\Delta G^B_{BL} = \beta^{-1} \ln \left[ \langle e^{-\beta u_{BL}(\mathbf{x})} \rangle^B_{\{BR,BA,\Theta,\Phi,\Psi,\phi,\theta\}} \right] \tag{10}$$
$$\Delta G^B_{\theta} = \beta^{-1} \ln \left[ \langle e^{-\beta u_{\theta}(\mathbf{x})} \rangle^B_{\{BR,BA,\Theta,\Phi,\Psi,\phi\}} \right] \tag{11}$$
$$\Delta G^B_{\phi} = \beta^{-1} \ln \left[ \langle e^{-\beta u_{\phi}(\mathbf{x})} \rangle^B_{\{BR,BA,\Theta,\Phi,\Psi\}} \right] \tag{12}$$
$$\Delta G^B_{\Psi} = \beta^{-1} \ln \left[ \langle e^{-\beta u_{\Psi}(\mathbf{x})} \rangle^B_{\{BR,BA,\Theta,\Phi\}} \right] \tag{13}$$
$$\Delta G^B_{\Phi} = \beta^{-1} \ln \left[ \langle e^{-\beta u_{\Phi}(\mathbf{x})} \rangle^B_{\{BR,BA,\Theta\}} \right] \tag{14}$$
$$\Delta G^B_{\Theta} = \beta^{-1} \ln \left[ \langle e^{-\beta u_{\Theta}(\mathbf{x})} \rangle^B_{\{BR,BA\}} \right] \tag{15}$$
$$\Delta G^B_{BA} = \beta^{-1} \ln \left[ \langle e^{-\beta u_{BA}(\mathbf{x})} \rangle^B_{\{BR\}} \right] \tag{16}$$
$$\Delta G^B_{BR} = \beta^{-1} \ln \left[ \langle e^{-\beta u_{BR}(\mathbf{x})} \rangle^B \right] \tag{17}$$
$$\Delta G^S_0 = \Delta G^S_\Theta + \Delta G^S_\Phi + \Delta G^S_\Psi + \Delta G^S_\theta + \Delta G^S_\phi \tag{18}$$
$$\Delta G^B_0 = \Delta G^B_\Theta + \Delta G^B_\Phi + \Delta G^B_\Psi = -\beta^{-1} \ln \left[ \frac{1}{8\pi^2} \int_0^\pi \int_0^{2\pi} \int_0^{2\pi} \sin(\Theta) e^{-\beta u_0(\Theta,\Phi,\Psi)} d\Psi d\Phi d\Theta \right] \tag{19}$$
$$I^* = \int_{bound} e^{-\beta[W(r)-W(r^*)]} dr \tag{20}$$
$$O^* = (r^*)^2 \int_0^\pi \int_0^{2\pi} \sin(\theta) e^{-\beta u_a(\theta,\phi)} d\theta d\phi \tag{21}$$
$$K_A^{Apo} = O^* I^* e^{-\beta[(\Delta G^B_{BR} - \Delta G^S_{BR}) + (\Delta G^B_{BA} - \Delta G^S_{BA}) + (\Delta G^B_0 - \Delta G^S_0)]} \tag{22}$$
$$K_A^{BL} = O^* I^* e^{-\beta[(\Delta G^B_{BR} - \Delta G^S_{BR}) + (\Delta G^B_{BA} - \Delta G^S_{BA}) + (\Delta G^B_0 - \Delta G^S_0) + (\Delta G^B_{BL} - \Delta G^S_{BL})]} \tag{23}$$
$$\Delta G^\circ = -\beta^{-1} \ln \left[ K_A C^\circ \right], \; C^\circ = \frac{1}{1661 \text{Å}^3} \tag{24}$$

Note that the $O^*$ and $\Delta G^B_0$ terms can be calculated analytically, as shown below, while each other term requires MD simulation.

**Apo orientational restraint contribution**

$$\begin{aligned}
O^*_{Apo} &= (r^*)^2 \int_0^{\pi} \int_0^{2\pi} \sin(\Theta) e^{-\beta u_a(\Theta,\Phi)} d\Phi d\Theta \\
&= (44\ \text{Å})^2 \int_0^{2\pi} e^{-\beta(.5)(.1)(180/\pi)^2(\Phi-16.8963(\pi/180))^2} d\Phi \int_0^{\pi} \sin(\Theta) e^{-\beta(.5)(.1)(180/\pi)^2(\Theta-120.4496(\pi/180))^2} d\Theta \\
&= (1936\ \text{Å}^2)(0.106819)(0.0920026) = 19.0262\ \text{Å}^2
\end{aligned}$$

**Apo bulk angular restraint contributions**

$$\begin{aligned}
\Delta G^B_{0,Apo} &= -\beta^{-1} \ln \left[ \frac{1}{8\pi^2} \int_0^{\pi} \int_0^{2\pi} \int_0^{2\pi} \sin(\Theta) e^{-\beta u_0(\Theta,\Phi,\Psi)} d\Psi d\Phi d\Theta \right] \\
&= -\beta^{-1} \ln \left[ \frac{1}{8\pi^2} \int_0^{2\pi} e^{-\beta(.5)(.1)(180/\pi)^2(\Psi-33.0273(\pi/180))^2} d\Psi \int_0^{2\pi} e^{-\beta(.5)(.1)(180/\pi)^2(\phi-261.461(\pi/180))^2} d\phi \right. \\
&\left. \int_0^{\pi} \sin(\theta) e^{-\beta(.5)(.1)(180/\pi)^2(\theta-73.4478(\pi/180))^2} d\theta \right] \\
&= 6.63\ \text{kcal} \cdot \text{mol}^{-1}
\end{aligned}$$

**Holo orientational restraint contribution**

$$\begin{aligned}
O^*_{Holo} &= (r^*)^2 \int_0^{\pi} \int_0^{2\pi} \sin(\Theta) e^{-\beta u_a(\Theta,\Phi)} d\Phi d\Theta \\
&= (44\ \text{Å})^2 \int_0^{2\pi} e^{-\beta(.5)(.1)(180/\pi)^2(\Phi-15.9178(\pi/180))^2} d\Phi \int_0^{\pi} \sin(\Theta) e^{-\beta(.5)(.1)(180/\pi)^2(\Theta-118.5843(\pi/180))^2} d\Theta \\
&= (1936\ \text{Å}^2)(0.106819)(0.0937143) = 19.3802\ \text{Å}^2
\end{aligned}$$

**Holo bulk angular restraint contributions**

$$\begin{aligned}
\Delta G^B_{0,Holo} &= -\beta^{-1} \ln \left[ \frac{1}{8\pi^2} \int_0^{\pi} \int_0^{2\pi} \int_0^{2\pi} \sin(\Theta) e^{-\beta u_0(\Theta,\Phi,\Psi)} d\Psi d\Phi d\Theta \right] \\
&= -\beta^{-1} \ln \left[ \frac{1}{8\pi^2} \int_0^{2\pi} e^{-\beta(.5)(.1)(180/\pi)^2(\Psi-33.7587(\pi/180))^2} d\Psi \int_0^{2\pi} e^{-\beta(.5)(.1)(180/\pi)^2(\phi-267.2762(\pi/180))^2} d\phi \right. \\
&\left. \int_0^{\pi} \sin(\theta) e^{-\beta(.5)(.1)(180/\pi)^2(\theta-75.9999(\pi/180))^2} d\theta \right] \\
&= 6.62\ \text{kcal} \cdot \text{mol}^{-1}
\end{aligned}$$

**Table S1:** Force constants and reference collective variable values used for restraints in REUS PMF calculations.

| Collective variable | $k_{Force}$ | Apo reference | Holo reference |
|---|---|---|---|
| BRI1 RMSD | 10.0* | 1.1825 Å | 1.1492 Å |
| BAK1 RMSD | 10.0* | 1.1973 Å | 1.3730 Å |
| $\Theta$ | $0.10^{\dagger}$ | 120.4496° | 118.5843° |
| $\Phi$ | $0.10^{\dagger}$ | 16.8963° | 15.9178° |
| $\Psi$ | $0.10^{\dagger}$ | 33.0273° | 33.7587° |
| $\phi$ | $0.10^{\dagger}$ | 261.461° | 267.2762° |
| $\theta$ | $0.10^{\dagger}$ | 73.4478° | 75.9999° |
| BL-BRI1 distance | 10.0* | N/A | 18.7735 Å |

$^{*}$kcal·mol$^{-1}$·Å$^{-2}$ $^{\dagger}$kcal·mol$^{-1}$·degree$^{-2}$

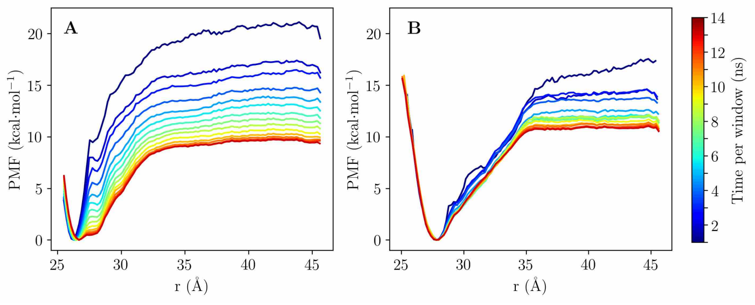


**Figure S1:** Convergence of the **A** apo and **B** holo tBRI1-BAK1 separation PMFs with REUS sampling time per window. This figure was produced using Matplotlib 2.2.2 [2].

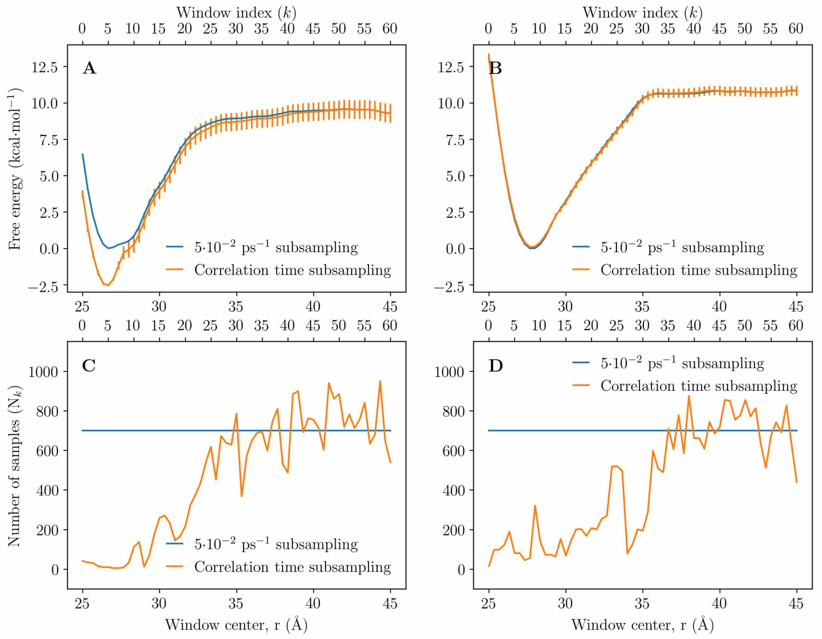


**Figure S2:** Rationale for our choice of a uniform 50 ns$^{-1}$ subsampling rate for REUS simulations. **A** The apo and **B** holo separation PMFs using a uniform 50 ns$^{-1}$ subsampling rate and a subsampling rate determined individually for each window using the correlation time method implemented in pymbar [3, 4]. The number of samples remaining for each window for the **C** apo and **D** holo systems. Using the correlation time to subsample yields uncorrelated samples but at the same time causes highly uneven sampling across $r$. Using a uniform subsampling time of 50 ns$^{-1}$ by definition yields even sampling over $r$, but likely results in the use of correlated samples in some windows, leading to underestimation of error [5]. This figure was produced using Matplotlib 2.2.2 [2].

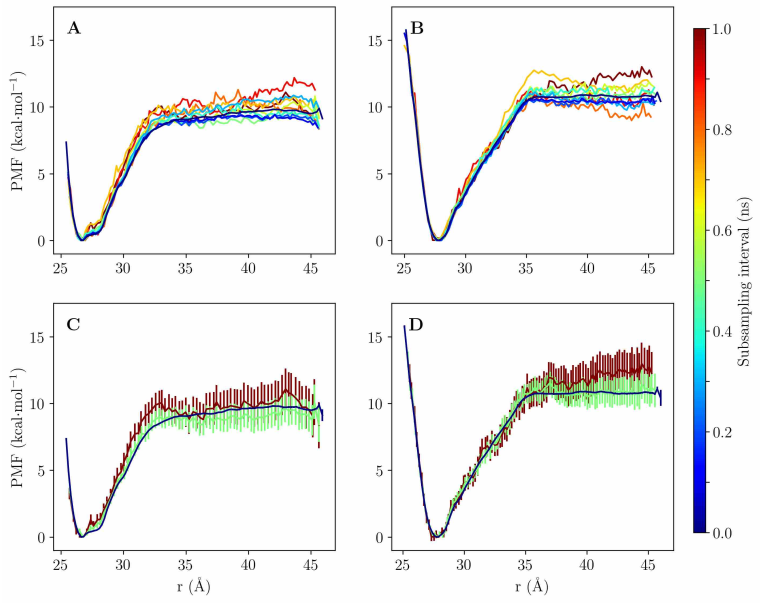


**Figure S3:** The effects of subsampling rate on the **A** apo and **B** holo separation PMFs. We chose three subsampling rates to include with error bars shown for the **C** apo and **D** holo separation PMFs. Note the general insensitivity to both separation PMFs to the choice of uniform subsampling rate over all windows. This figure was produced using Matplotlib 2.2.2 [2].

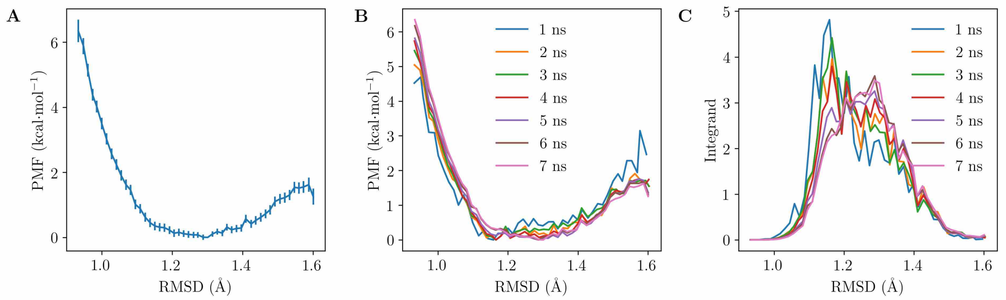


**Figure S4:** tBRI1 conformational restraint contributions for bound apo BRI1-BAK1. **A** The restraint PMF. **B** Convergence of the restraint PMF with simulation time. **C** Convergence of the integrand within the associated ensemble average estimation with time. This figure was produced using Matplotlib 2.2.2 [2].

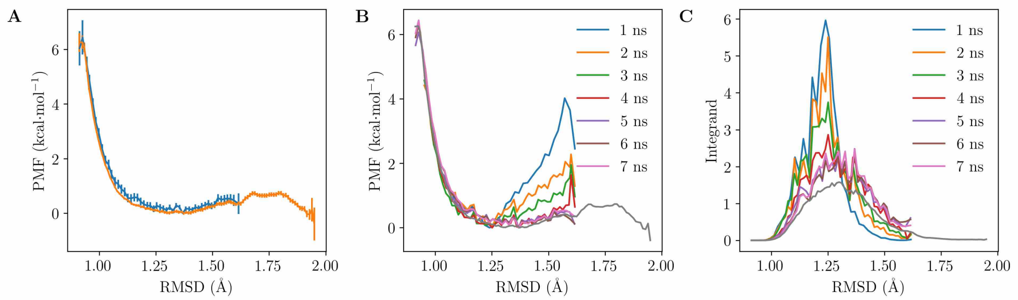


**Figure S5:** BAK1 conformational restraint contributions for bound apo BRI1-BAK1. **A** The restraint PMF. The PMF including additional umbrella sampling is shown in orange. **B** Convergence of the restraint PMF with simulation time. The PMF including additional umbrella sampling is shown in grey. **C** Convergence of the integrand within the associated ensemble average estimation with time. The integrand including additional umbrella sampling is shown in orange. This figure was produced using Matplotlib 2.2.2 [2].

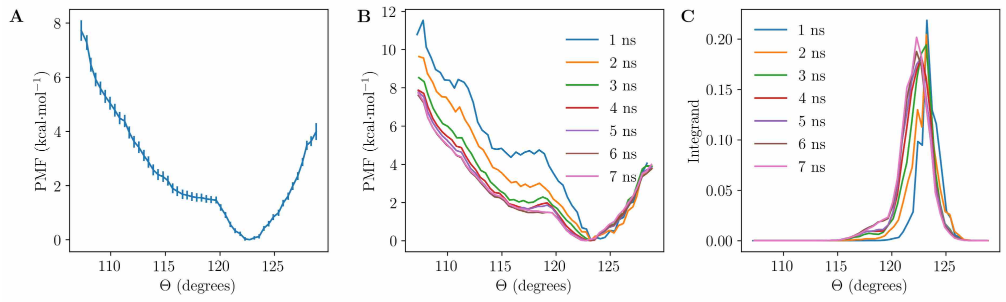


**Figure S6:** $\Theta$ restraint contributions for bound apo BRI1-BAK1. **A** The restraint PMF. **B** Convergence of the restraint PMF with simulation time. **C** Convergence of the integrand within the associated ensemble average estimation with time. This figure was produced using Matplotlib 2.2.2 [2].

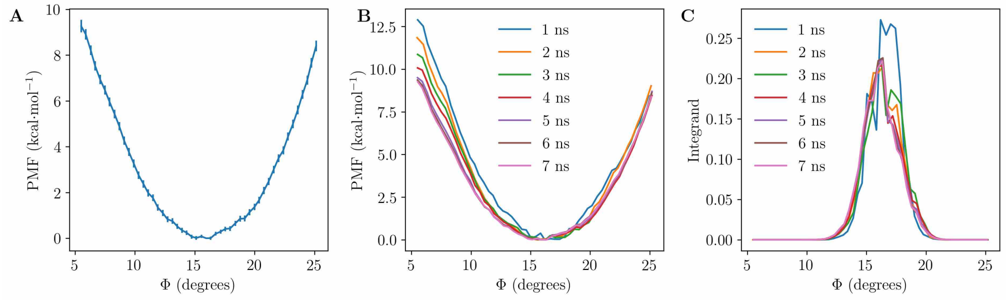


**Figure S7:** $\Phi$ restraint contributions for bound apo BRI1-BAK1. **A** The restraint PMF. **B** Convergence of the restraint PMF with simulation time. **C** Convergence of the integrand within the associated ensemble average estimation with time. This figure was produced using Matplotlib 2.2.2 [2].

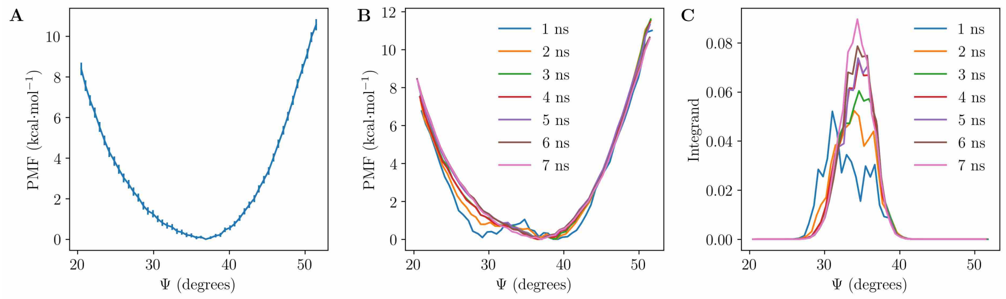


**Figure S8:** $\Psi$ restraint contributions for bound apo BRI1-BAK1. **A** The restraint PMF. **B** Convergence of the restraint PMF with simulation time. **C** Convergence of the integrand within the associated ensemble average estimation with time. This figure was produced using Matplotlib 2.2.2 [2].

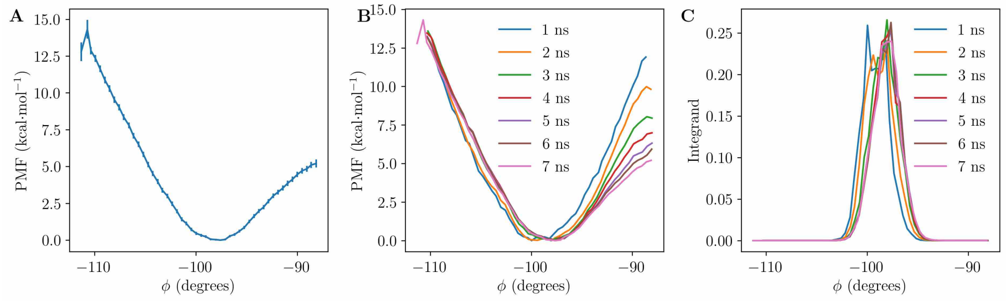


**Figure S9:** $\phi$ restraint contributions for bound apo BRI1-BAK1. **A** The restraint PMF. **B** Convergence of the restraint PMF with simulation time. **C** Convergence of the integrand within the associated ensemble average estimation with time. This figure was produced using Matplotlib 2.2.2 [2].

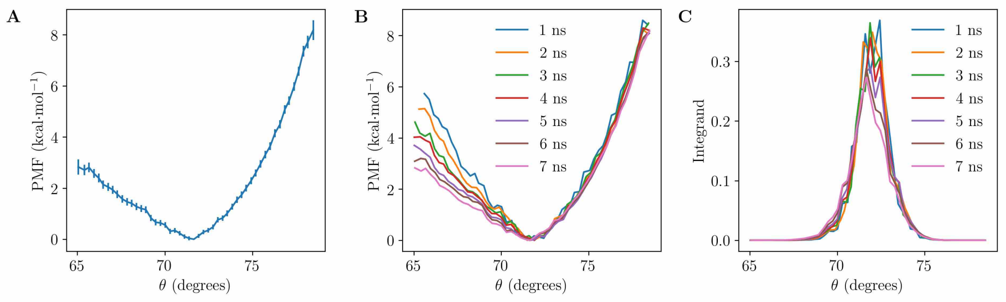


**Figure S10:** $\theta$ restraint contributions for bound apo BRI1-BAK1. **A** The restraint PMF. **B** Convergence of the restraint PMF with simulation time. **C** Convergence of the integrand within the associated ensemble average estimation with time. This figure was produced using Matplotlib 2.2.2 [2].

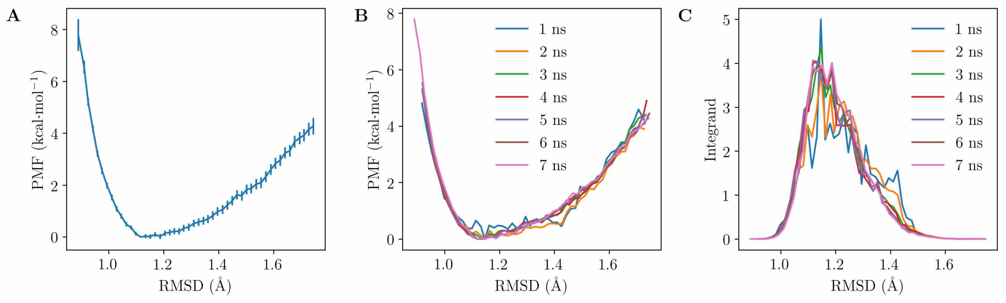


**Figure S11:** BAK1 conformational restraint contributions for unbound apo BRI1-BAK1. **A** The restraint PMF. **B** Convergence of the restraint PMF with simulation time. **C** Convergence of the integrand within the associated ensemble average estimation with time. This figure was produced using Matplotlib 2.2.2 [2].

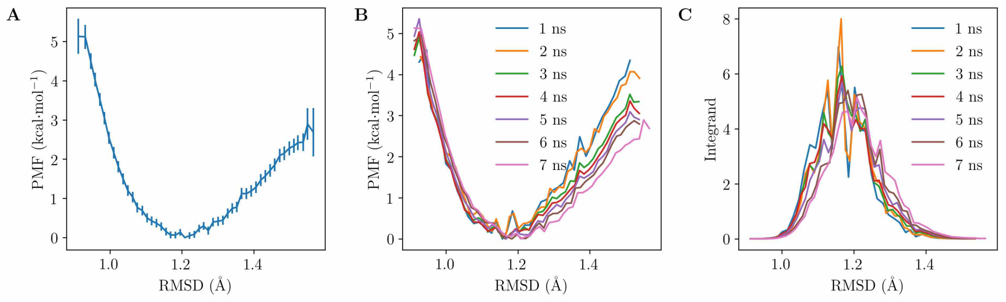


**Figure S12:** tBRI1 conformational restraint contributions for unbound apo BRI1-BAK1. **A** The restraint PMF. **B** Convergence of the restraint PMF with simulation time. **C** Convergence of the integrand within the associated ensemble average estimation with time. This figure was produced using Matplotlib 2.2.2 [2].

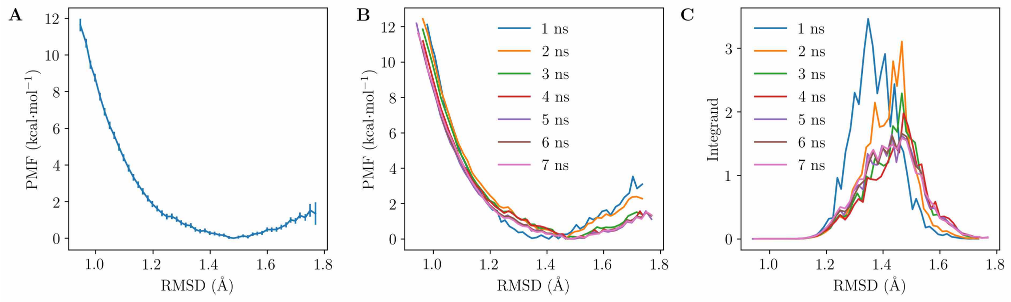


**Figure S13:** tBRI1 conformational restraint contributions for bound holo BRI1-BAK1. **A** The restraint PMF. **B** Convergence of the restraint PMF with simulation time. **C** Convergence of the integrand within the associated ensemble average estimation with time. This figure was produced using Matplotlib 2.2.2 [2].

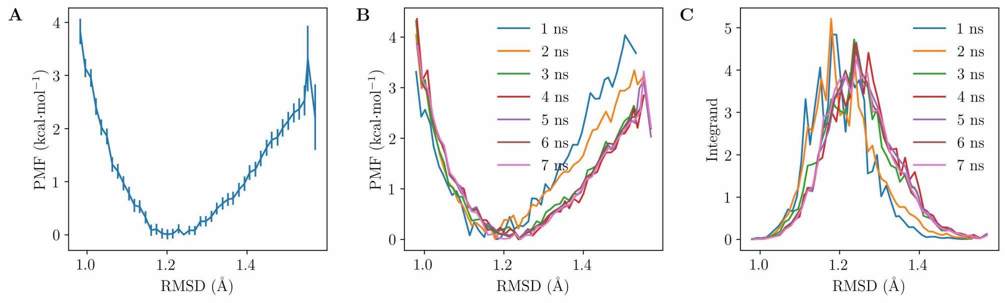


**Figure S14:** BAK1 conformational restraint contributions for bound holo BRI1-BAK1. **A** The restraint PMF. **B** Convergence of the restraint PMF with simulation time. **C** Convergence of the integrand within the associated ensemble average estimation with time. This figure was produced using Matplotlib 2.2.2 [2].

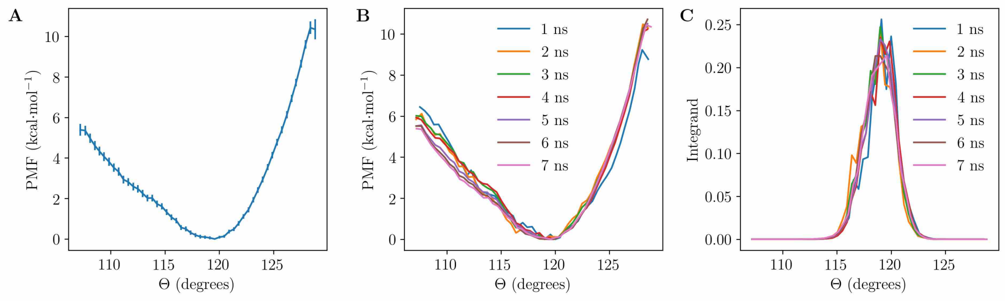


**Figure S15:** Θ restraint contributions for bound holo BRI1-BAK1. **A** The restraint PMF. **B** Convergence of the restraint PMF with simulation time. **C** Convergence of the integrand within the associated ensemble average estimation with time. This figure was produced using Matplotlib 2.2.2 [2].

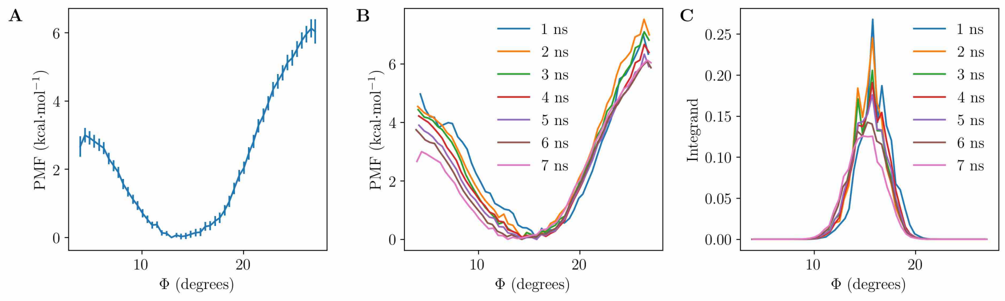


**Figure S16:** Φ restraint contributions for bound holo BRI1-BAK1. **A** The restraint PMF. **B** Convergence of the restraint PMF with simulation time. **C** Convergence of the integrand within the associated ensemble average estimation with time. This figure was produced using Matplotlib 2.2.2 [2].

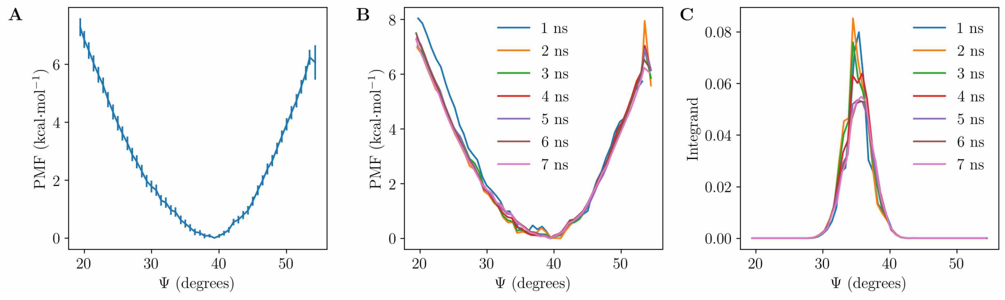


**Figure S17:** Ψ restraint contributions for bound holo BRI1-BAK1. **A** The restraint PMF. **B** Convergence of the restraint PMF with simulation time. **C** Convergence of the integrand within the associated ensemble average estimation with time. This figure was produced using Matplotlib 2.2.2 [2].

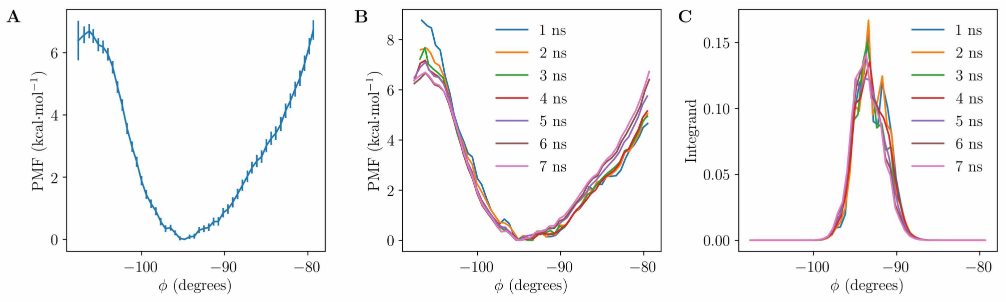


**Figure S18:** $\phi$ restraint contributions for bound holo BRI1-BAK1. **A** The restraint PMF. **B** Convergence of the restraint PMF with simulation time. **C** Convergence of the integrand within the associated ensemble average estimation with time. This figure was produced using Matplotlib 2.2.2 [2].

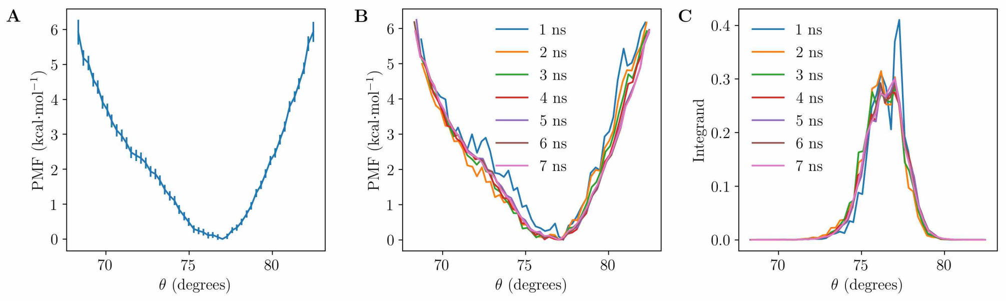


**Figure S19:** $\theta$ restraint contributions for bound holo BRI1-BAK1. **A** The restraint PMF. **B** Convergence of the restraint PMF with simulation time. **C** Convergence of the integrand within the associated ensemble average estimation with time. This figure was produced using Matplotlib 2.2.2 [2].

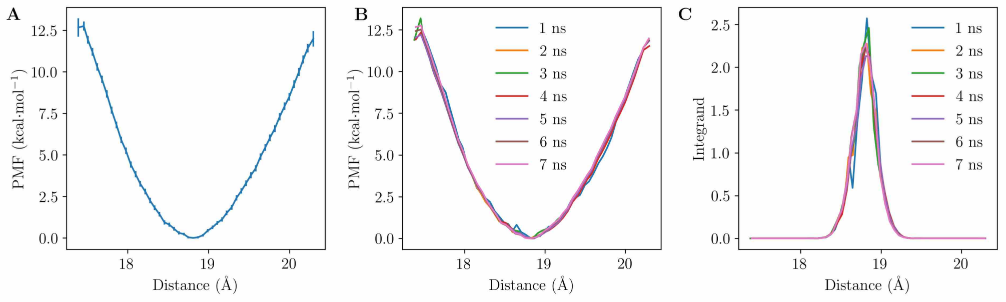


**Figure S20:** BL restraint contributions for bound holo BRI1-BAK1. **A** The restraint PMF. **B** Convergence of the restraint PMF with simulation time. **C** Convergence of the integrand within the associated ensemble average estimation with time. This figure was produced using Matplotlib 2.2.2 [2].

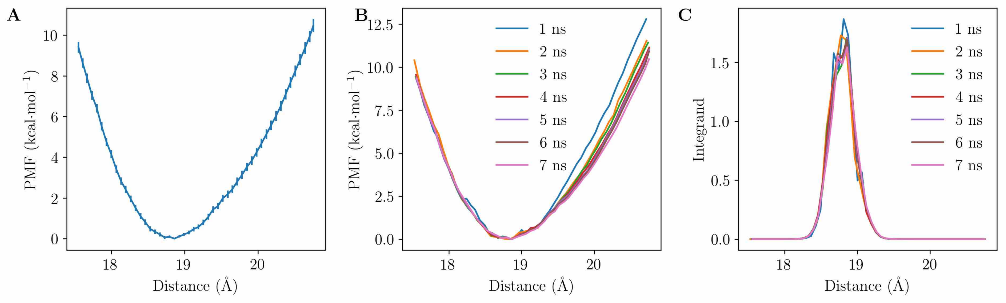


**Figure S21:** BL restraint contributions for unbound holo BRI1-BAK1. **A** The restraint PMF. **B** Convergence of the restraint PMF with simulation time. **C** Convergence of the integrand within the associated ensemble average estimation with time. This figure was produced using Matplotlib 2.2.2 [2].

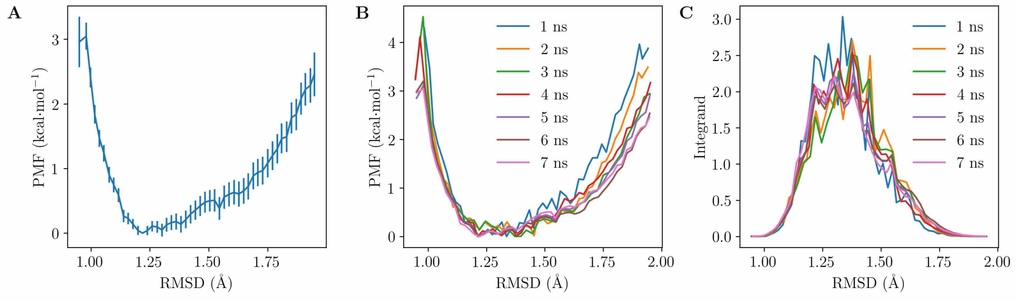


**Figure S22:** BAK1 conformational restraint contributions for unbound holo BRI1-BAK1. **A** The restraint PMF. **B** Convergence of the restraint PMF with simulation time. **C** Convergence of the integrand within the associated ensemble average estimation with time. This figure was produced using Matplotlib 2.2.2 [2].

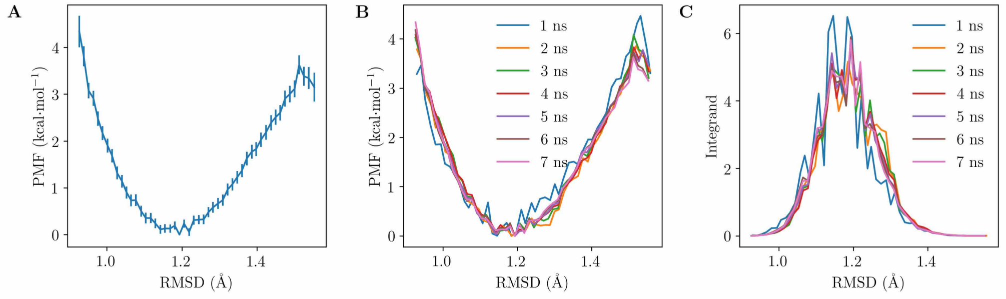


**Figure S23:** tBRI1 conformational restraint contributions for unbound holo BRI1-BAK1. **A** The restraint PMF. **B** Convergence of the restraint PMF with simulation time. **C** Convergence of the integrand within the associated ensemble average estimation with time. This figure was produced using Matplotlib 2.2.2 [2].

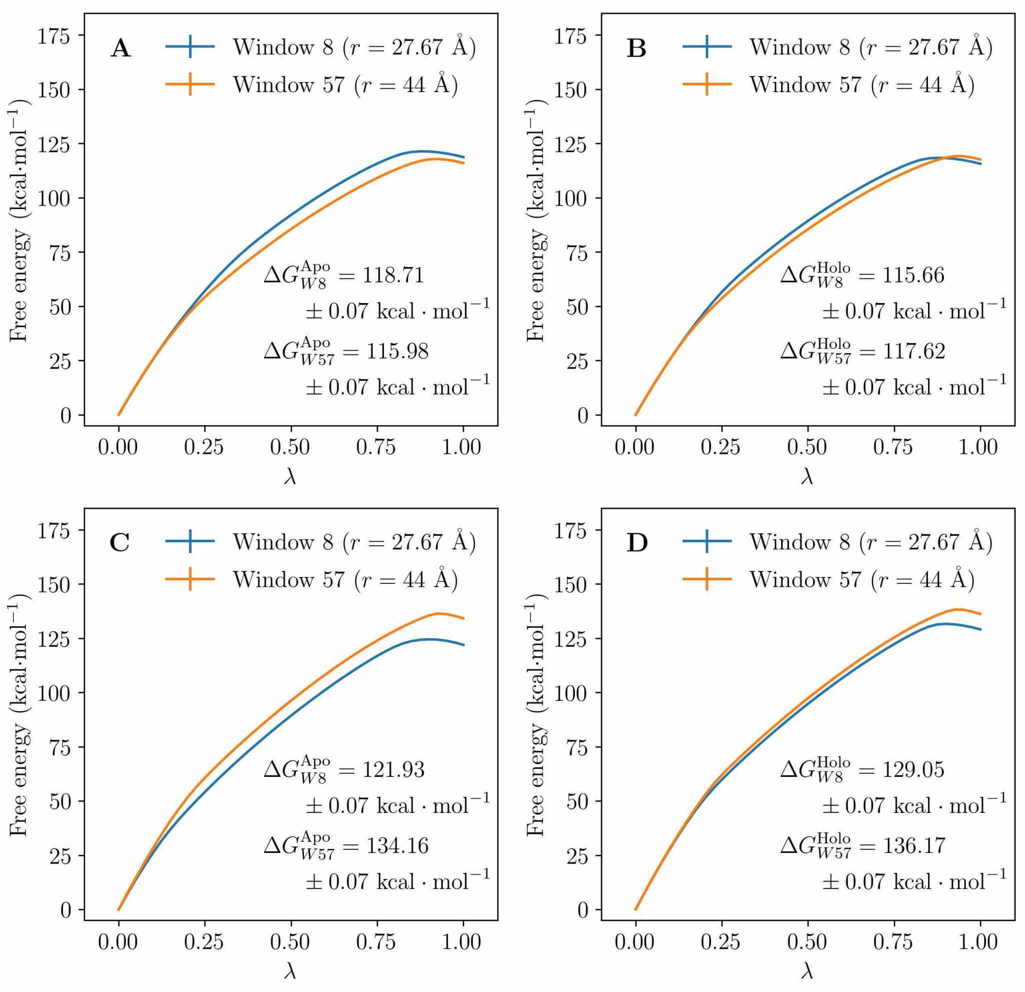


**Figure S24:** Results from alchemical free energy calculations describing protonation of BAK1 H61 and BRI1 E749, each performed with BRI1 and BAK1 close (window 8, $r = 27.67$ Å) and distant (window 57, $r = 44.00$ Å). On the x-axis is the alchemical parameter $\lambda$ while on the y-axis is the MBAR-derived relative free energy of each state. **A** Protonation of apo BAK1 H61. **B** Protonation of holo BAK1 H61. **C** Protonation of apo BRI1 E749. **D** Protonation of holo BRI1 E749. This figure was produced using Matplotlib 2.2.2 [2].

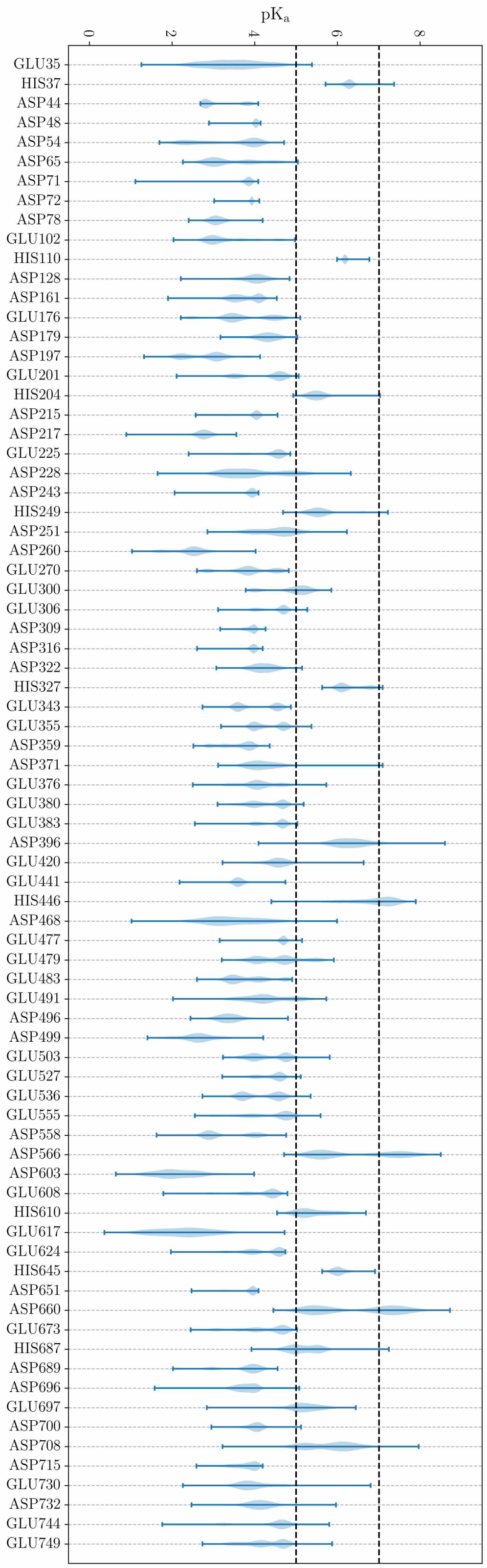


**Figure S25:** Violin plot of BRI1 sidechain $pK_a$s for residues with sidechain $pK_a$s close to 5, calculated from the simulations of the full, apo BRI1 ECD using PROPKA 3.1 [6, 7]. Frames were taken from apo BRI1 simulations at a rate of 10 $ns^{-1}$. Bars represent the interval from the lowest to the highest calculated $pK_a$ over all frames. This figure was produced using Matplotlib 2.2.2 [2].
